# *Discoidin domain receptor* regulates ensheathment, survival, and caliber of peripheral axons

**DOI:** 10.1101/2021.09.21.461270

**Authors:** Megan M. Corty, Alexandria P. Lassetter, Jo Q. Hill, Amy E. Sheehan, Sue A. Aicher, Marc R. Freeman

## Abstract

Invertebrate axons and small caliber axons in mammalian peripheral nerves are unmyelinated but still ensheathed by glia. How this type of ensheathment is controlled and its roles in supporting neuronal function remain unclear. Here we use *Drosophila* wrapping glia, which ensheathe peripheral axons to study the function and development of non-myelinating ensheathment. We developed a new SplitGal4 intersectional driver to target wrapping glia for genetic ablation and found that loss of wrapping glia severely impaired larval locomotor behavior. We performed an *in vivo* RNAi screen in *Drosophila* to identify glial genes required for axon ensheathment during development and identified the conserved receptor tyrosine kinase Discoidin domain receptor (Ddr). In larval peripheral nerves, loss of Ddr resulted in severely reduced ensheathment of axons. We found a strong dominant genetic interaction between *Ddr* and the fly type XV/XVIII collagen *Multiplexin (Mp)*, suggesting Ddr functions a collagen receptor to drive wrapping of axons during development. Surprisingly, despite severe impairment of ensheathment, the residual wrapping in *Ddr* mutants was sufficient to support basic circuit function during larval stages. In adult nerves, loss of Ddr from glia decreased long-term survival of sensory neurons and significantly reduced axon caliber in an identifiable neuron without overtly affecting ensheathment. Our data establish a crucial role for non-myelinating glia in peripheral nerve development and function across the lifespan, and identify Ddr as a key regulator of axon-glia interactions during ensheathment and nerve growth.

## Introduction

In complex nervous systems, specialized glial cells ensheathe long axons, and this wrapping can take many forms. Myelination is the most studied type of ensheathment, but unmyelinated axons make up the majority (∼70%) of axons in human peripheral nerves (Schmalbruch 1986; Ochoa and Mair 1969). Vertebrate Remak Schwann cells ensheathe and separate unmyelinated axons from one another. These small caliber axons include many types of sensory neuron axons, including nociceptive c-fibers and autonomic axons (Griffin and Thompson 2008). Remak Schwann cells are positioned to mediate fundamental aspects of nerve development and sensory biology, and be important modulators of neurological conditions affecting the peripheral nervous system (PNS) including peripheral neuropathies and nerve injuries. However, Remak Schwann cells have remained understudied, in part due to a lack of selective genetic tools to specifically target this population of cells.

In *Drosophila,* axons in peripheral nerves are ensheathed by specialized wrapping glia in a manner analogous to vertebrate Remak bundles (Figure 1A). *Drosophila* larval abdominal nerves project out of the ventral nerve cord in each segment and contain both motor and sensory neuron axons surrounded by multiple glial layers: the outermost perineurial glia, then subperineural glia, and finally axon-associated wrapping glia (Stork et al. 2008; Matzat et al. 2015; Hilchen et al. 2013)(Figure 1A). Wrapping is progressive throughout larval life. In embryos, axons are initially tightly fasciculated without intervening glial processes. Wrapping progresses such that by the third larval instar (∼3-4 days later) axons are wrapped either individually or in small bundles by wrapping glia membrane (Figure 1B) (Matzat et al. 2015). This process mirrors vertebrate nerve development when Schwann cells perform radial sorting of axons that are initially tightly fasciculated (Monk et al. 2015) and a growing body of evidence that demonstrates a high degree of cellular and molecular conservation between ensheathment mechanisms in *Drosophila* and vertebrates (Matzat et al. 2015; Ghosh et al. 2013; Xie and Auld 2011; Petley-Ragan et al. 2016; Mukherjee et al. 2020). This Remak-like multi-axonal ensheathment is thought to represent an ancient form of axon-glial association, and the emerging cellular and molecular conservation of multiple aspects of peripheral nerve biology make *Drosophila* wrapping glia a promising model to study non-myelinating ensheathment *in vivo*.

**Figure 1:**
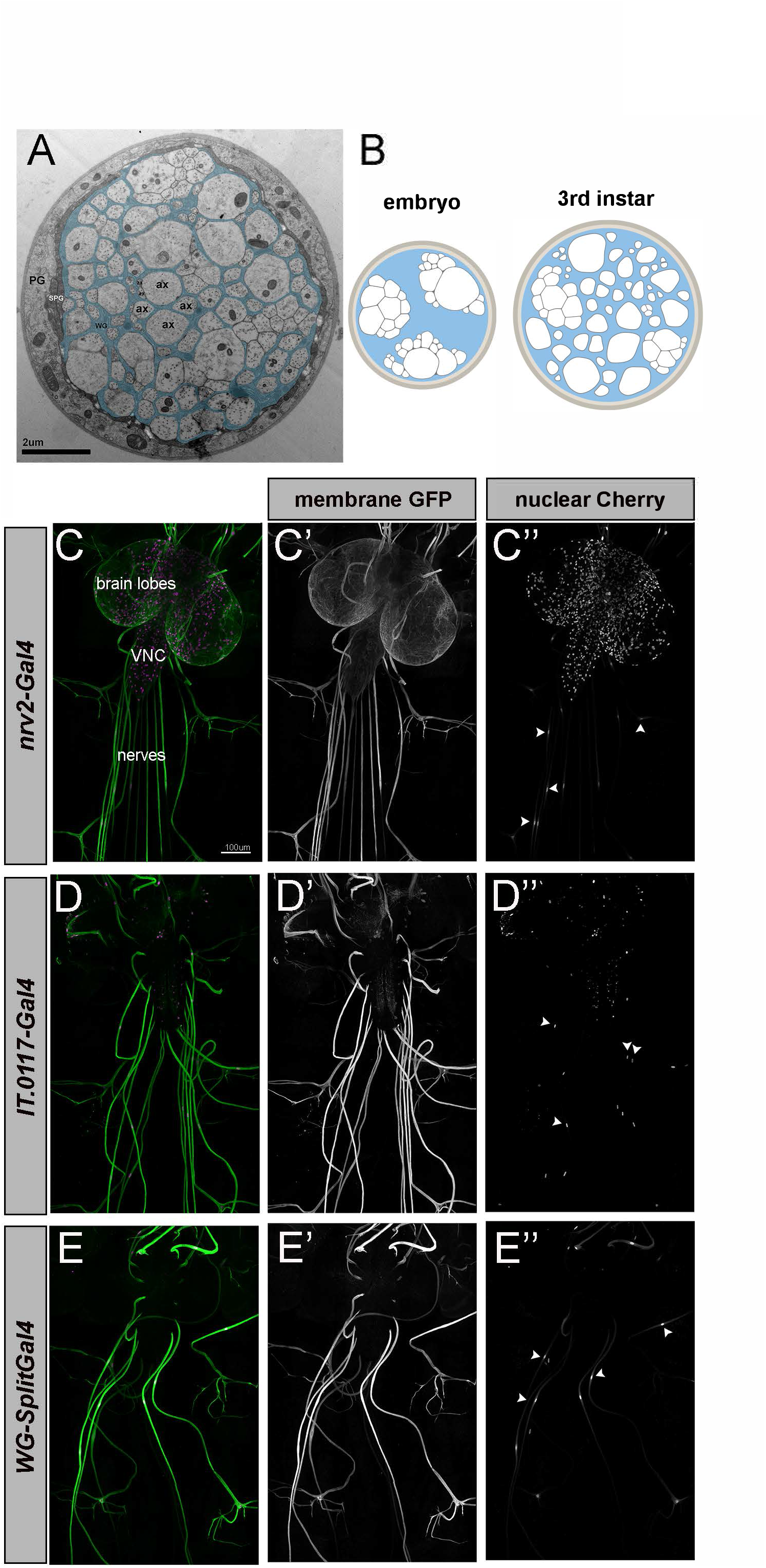
Construction of *Drosophila* wrapping glia *SplitGal4* driver. **(A)** TEM cross-section of a third instar larval nerve. Light, round profiles of larval axons (ax) are surrounded by darker (pseudo-colored cyan) wrapping glia. Subperineurial (SPG) and perineurial glia (PG; not colored) form the outer layers of the nerve, which is coated by a neural lamella of ECM proteins. **(B)** Schematic representation of larval nerve cross sections at embryo hatching and 3^rd^ instar stages illustrating that wrapping is progressive over larval development. **(C-D)** Confocal images showing 3^rd^ instar larval nervous systems including the brain, ventral nerve cord (VNC), and peripheral nerves. **(C)** Expression pattern of *nrv2-Gal4* driving *UAS-CD8:GFP* (green, C’) and *UAS-mCherry^nls^* (magenta, C’’). *Nrv2-Gal4* drives exclusively in wrapping glia in the periphery and three CNS glia subtypes--cortex glia, ensheathing glia, and astrocytes as can be seen with the labeling throughout the brain and VNC. **(D)** Expression pattern of *IT.0117-Gal4* driving *UAS-myrGFP* (green, D’) and *UAS-H2B:mCherry* (magenta, D’’). *IT.0117-Gal4* drives in wrapping glia in the periphery and a subset of CNS neurons that appear to be primarily interneurons. **(E)** Expression pattern of *Wrapping glia-SplitGal4* (made by combining *nrv2-Gal4^DBD^* and *IT.0117-Gal4^VP16AD^*) driving *UAS-mCD8:GFP* (green, E’) and *UAS-mCherry^nls^* (magenta, E’’). *WG-SplitGal4* drives exclusively in wrapping glia in the periphery without any evidence of neuronal or glial expression in the CNS. Arrowheads indicate a subset of wrapping glia nuclei in each panel.

Beyond simply insulating individual axons, glia that ensheathe axons also regulate neuronal development, maintenance, and function throughout an animal’s lifespan. Myelination can alter the distribution of axonal proteins, increase axon caliber, and is important for trophic and metabolic support of long axons, though the precise mechanisms by which myelinating glia perform all of these functions remain incompletely understood (Nave 2010). Recent work supports the notion that Remak Schwann cells play similar important roles: perturbation of Schwann cell metabolism results in axon degeneration, with small caliber, Remak-ensheathed axons degenerating earlier than myelinated ones (Beirowski et al. 2014). Although Remak Schwann cells were not targeted selectively, these observations support the notion that unmyelinated axons are also reliant on glial support.

Here we take advantage of *Drosophila* wrapping glia as a model to identify new molecular regulators of glial ensheathment and support of axons. We demonstrate that wrapping glia are essential for normal nerve function and behavior by developing a new, highly precise Split-Gal4 intersectional driver to genetically ablate larval wrapping glia, which results in severely impaired crawling behavior. Through an RNAi-based screen, we identified the conserved Discoidin domain receptor (Ddr) as crucial for glial ensheathment in larval nerves. Surprisingly, while loss of Ddr resulted in incomplete wrapping, it did not result in overt effects on behavior, arguing that even partial ensheathment is sufficient to support normal nerve function. In adult nerves, loss of *Ddr* resulted in reduced long-term survival of peripheral sensory neurons and reduced axon caliber without affecting ensheathment. *Ddr* is therefore a new glial regulator of axon caliber *in vivo*.

Finally, we identify the collagen Multiplexin (Mp) as a potential glial-derived ligand for Ddr in driving wrapping during development, suggesting that Ddr-collagen interactions are critical regulators of axon ensheathment and glial support of neurons *in vivo*.

## RESULTS

### Genetic ablation of wrapping glia impairs larval crawling

To determine whether axonal ensheathment is required for normal axon conduction or circuit function, we sought to selectively ablate wrapping glia. *Nrv2-Gal4* is a Gal4 driver commonly used to manipulate wrapping glia because it drives strong expression of UAS transgenes specifically in wrapping glia in the PNS without any expression in neurons. Although *Nrv2-Gal4* is specific to wrapping glia in the PNS, it also drives expression in some subtypes of CNS glia which complicates the interpretation of ablation experiments (Figure 1C). To generate a more precise wrapping glia-specific driver line, we turned to a Split-Gal4 intersectional strategy (Luan et al. 2006). First, we created a *nrv2-Gal4^DBD^* construct. This line is capable of driving expression of the DNA binding domain of GAL4 in wrapping glia, cortex glia, ensheathing glia, and astrocytes (Coutinho-Budd et al. 2017). We then identified a line from the InSite collection (*InSite0117-Gal4*) that robustly labeled wrapping glia and a subset of neurons (Figure 1D). We converted *Insite0117*-*Gal4* to *Insite0117-GAL4^VP16AD^* line using the InSite method to swap the domains via genetic crosses (Gohl et al. 2011). In conjunction with *Nrv2-Gal4^DBD^* this results in specific expression of UAS-transgenes where expression of the 2 hemi-drivers overlaps—in wrapping glia. No other glia or neurons were labeled by this combination of drivers (Figure 1E and Supplemental Figure S1), which hereafter is referred to as *wrapping glia-Split Gal4* (*WG-SplitGal4*).

We used *WG-SplitGal4* to drive expression of *UAS-mCD8:GFP* and *UAS-reaper* to ablate wrapping glia. We observed nearly complete loss of larval wrapping glia based on loss of fluorescent reporters and an independent wrapping glia nuclear marker that we had identified in the lab, Oaz (Figure 2A-B and Supplemental Figures S2 & S3). To further confirm the ablation of wrapping glia, we examined the ultrastructure of nerves where wrapping glia were ablated by transmission electron microscopy (TEM). We saw tightly fasciculated axons without intervening glial membranes confirming wrapping glia had been eliminated (Figure 2C). Only in a few nerves did we identify anything that might be remnants of wrapping glia. We did not observe any obvious ingrowth of the outer subperineurial or perineurial glia layers between axons to potentially compensate for the loss of wrapping glia. However, several nerves showed an abnormal hypertrophy of the outer perineurial glia layer (Supplemental Figure S3). Interestingly, when we counted the number of axon profiles in each nerve, we found fewer axonal profiles than the expected ∼78 of A3-A7 nerves (*w^1118^* average= 77.4 axons; WG-ablated average= 63 axons; Unpaired t-test p=0.0006), indicating that axons (or entire neurons) might have died or not developed properly as a result of acute wrapping glia loss (Supplemental Figure S3).

**Figure 2:**
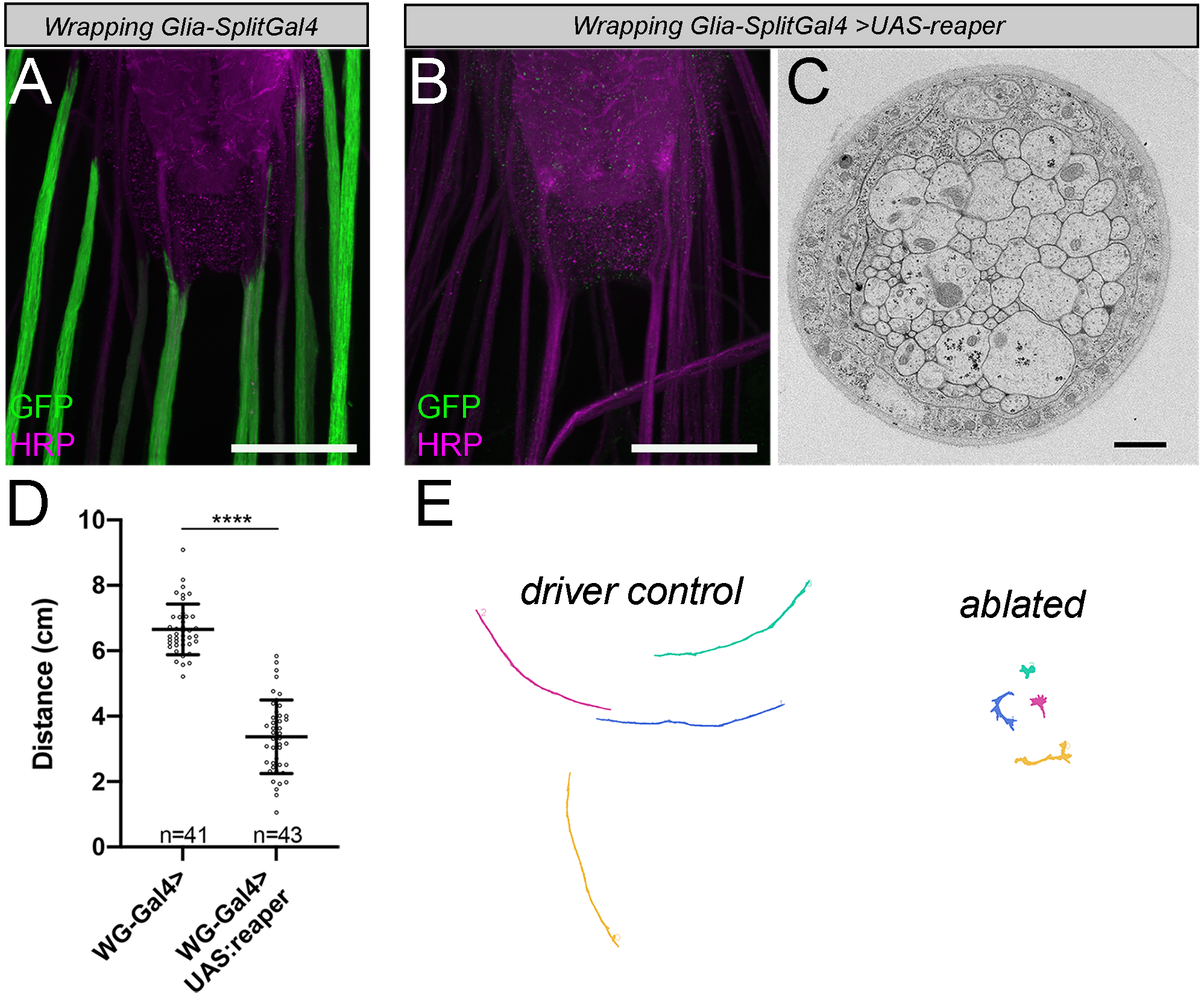
Genetic ablation of wrapping glia impairs larval crawling behavior. **(A)** Expression pattern of *WG-Split-Gal4* driving *UAS-CD8:GFP* (green). Magenta is an **HRP** counterstain which labels neuronal membranes. Scale bar, 50μm. **(B)** Representative image of a 3^rd^ instar larvae in which wrapping glia have been genetically ablated. Note the lack of GFP staining along the nerves. Scale bar 50μm. **(C)** TEM image of a nerve cross section from a larva whose wrapping glia have been ablated. All of the axons remain in contact with each other without intervening glial membrane. Scale bar 1 μm **(D)** Larval crawling behavior is impaired when wrapping glia are ablated. Unpaired t-test, p<0.0001. n = # of larvae/condition. **(E)** Representative crawling paths of control and ablated larvae. The paths from the ablated larvae are shorter and show increased turning, stalling, and reversals. Error bars: standard deviation

Using these tools, we assayed larval crawling behavior in *Ddr* mutant third instar larvae by monitoring crawling using the FIMTracker system (Risse et al. 2017). We found that animals with ablated wrapping glia crawled significantly less than control animals (Figure 2D-E; mean distance travelled: driver control = 6.65±0.77cm, n=41 larvae, ablated = 3.37±1.126cm, n=43; p<0.0001 unpaired t-test). We also observed that larvae lacking wrapping glia had difficulty righting themselves when placed on their dorsal sides, and exhibited abnormal postures and body bends, indicating a possible disruption of bilateral or intersegmental coordination caused by wrapping glia ablation. Together, these data demonstrate that wrapping glia are required in nerves for normal circuit function.

### Morphology based RNAi screen identifies *Ddr* as a regulator of wrapping glia development

To identify genes that are required in glia for normal ensheathment of axons, we conducted a morphology-based RNAi screen: we systematically knocked down genes in wrapping glia and assayed for changes in glial morphology in third instar larval nerves (Matzat et al. 2015; Stork et al. 2008). We used the well-established *nrv2-Gal4* driver line which, within the PNS, is strongly and exclusively expressed in wrapping glia, and appears much stronger than our *WG-SplitGal4* (Stork et al. 2008; Xie and Auld 2011). We expressed a membrane bound marker (*UAS-myr:tdTomato*) to visualize wrapping glia morphology and *UAS-RNAi* constructs. In total, we screened a collection of ∼2000 RNAi lines, which comprise the majority of transmembrane, secreted, and signaling proteins in the fly genome and included the fly homologs of ∼200 genes strongly expressed in the developing mouse oligodendrocyte lineage based on available RNA-seq data (Zhang et al. 2014).

Using spinning disk confocal microscopy and examination of nerve cross sections, we found that wild type wrapping glia morphology in third instar larvae includes membrane coverage throughout the interior of the nerve with small, regularly spaced gaps like a tight honeycomb (Figure 3B). As a positive control for our screen, we knocked down genes previously shown to be involved in wrapping glia development including the neuregulin homolog *vein*, the FGF receptor *heartless*, integrin receptors, laminin, and the ceramide synthetase gene, *schlank* (Matzat et al. 2015; Kottmeier et al. 2020; Ghosh et al. 2013; Xie and Auld 2011; Petley-Ragan et al. 2016; Franzdóttir et al. 2009), all of which resulted in easily identifiable defects in axon ensheathment using confocal microscopy (Supplemental Figure S4). Each of these genes have homologs that have been implicated in oligodendrocyte and/or Schwann cell development, further arguing for strong molecular conservation among both vertebrate and invertebrate axonal ensheathment mechanisms (Feltri et al. 2002; Barros et al. 2009; Pereira et al. 2009; Taveggia et al. 2005; Michailov et al. 2004; Stassart et al. 2013; Lyons et al. 2005; Yu et al. 2009; Furusho et al. 2009).

**Figure 3:**
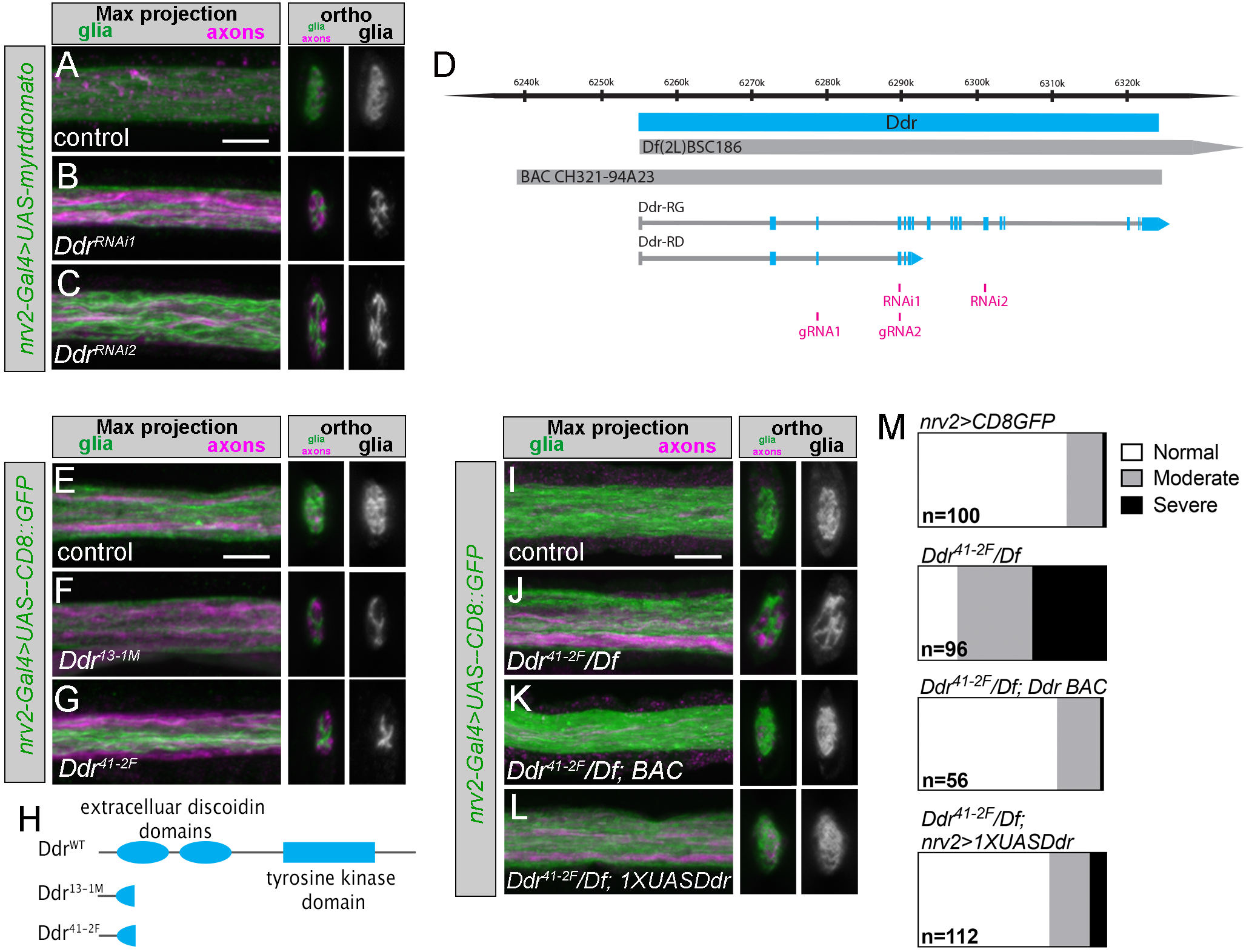
Ddr is required for normal wrapping glia morphogenesis. (**A-C**) Representative images ofDdr RNAi knockdown with 2 independent RNAi constructs. *Nrv2-Ga14* driven tdTomato is pseudocolored green. A subset ofaxons is labeled with anti-Futsch antibody (magenta). Compared to control nerves (**A**), RNAi conditions show large patches ofthe nerve cross section without glia membrane coverage (**B,C**). (**D**) Model ofthe *Ddr* genetic locus showing the coding region (cyan); the deficiency used in experiments: *Df(2L)BSC186;* the region that the rescue BAC covers; the location ofthe RNAi target regions (magenta); and locations ofthe guide RNAs used for the CRISPRmediated mutagenesis (magenta). (**E-G**) Representative images of *Ddr* homozygous mutant experiments. Compared to control (E) both CRISPR mutant alleles show severe defective glia coverage in the nerve cross sections (**F-G**). (**H**) Ddr is a transmembrane receptor tyrosine kinase characterized by extracellular discoidin and discoidin-like domains and an intracellular tyrosine kinase domain. The mutant alleles generated by CRlSPR-CAS9 result in a severely truncated peptide without any full domains. (**I-L**) Representative images of *Ddr* loss of function experiments. (**I**) *Nrv2-Gal4, UAS-CD8:GFP/+* was used to visualize glial morphology and is in the background of all conditions. (**J**) *Ddr^41-2F^IDjBSC186 (“Ddr* mutant”) nerves show incomplete coverage of the nerve cross section. This example shows a "moderate" phenotype, more examples of moderate and severe phenotypes can be seen in Supplemental Figure S2A. (K) Introduction of a BAC transgene to resupply endogenous *Ddr* expression largely restores normal morphology. (**L**) Expression of a *lxUAS-Ddr* construct in wrapping glia using *Nrv2-Ga14* similarly restores normal morphology. (**M**) Categorical scoring of nerve wrapping glia phenotypes. n= # of nerves scored (8-14 larvae per condition). Scale bars = 5μm.

Among potential hits whose function had not previously been studied in either vertebrate or invertebrate glia was the *Discoidin domain receptor* (*Ddr*) gene. Knocking down *Ddr* using two independent, non-overlapping RNAi constructs led to altered wrapping glia morphology with the nerve cross section incompletely covered by glial membrane (Figure 3A-C). *Drosophila Ddr* encodes a receptor tyrosine kinase homologous to vertebrate Ddr1 and Ddr2. In the mouse, Ddr1 is expressed in Schwann cells and the oligodendrocyte lineage (Zhang et al. 2014; Gerber et al. 2021; Roig et al. 2010; Franco-Pons et al. 2006), but direct roles for Ddr1in regulating ensheathment of axons have not been reported. Our data provide the first functional evidence that this receptor regulates axonal ensheathment.

### *Ddr* mutants exhibit defects in axonal ensheathment

To confirm that *Ddr* is involved in wrapping glia development, we created mutant alleles using a CRISPR-Cas9 strategy. Briefly, using two gRNAs against widely spaced, adjacent exons, we generated fly stocks in which an ∼11kb region of the *Ddr* coding region was excised near the 5’ end (Figure 3D). *Ddr* encodes two potential isoforms: a short isoform that is 380 amino acids and lacks both transmembrane and kinase domains, and a full-length isoform that encodes a 1054 amino acid protein. We identified two independent alleles that are predicted nulls, as they result in 150 amino acid (*Ddr^13-1M^*) or 134 amino acid (*Ddr^41-2F^*) long peptides lacking all known functional domains (Figure 3H). Both mutant alleles were homozygous viable and viable when placed in *trans* to large deficiencies that uncover the *Ddr* coding region.

We analyzed the nerves of *Ddr* homozygous mutant animals using florescent confocal microscopy and found that wrapping glia morphology was impaired (Figure 3E-G). We focused our loss of function analyses on using the combination of *Ddr^41-2F^* and *Df(2L) BSC186*, the shortest deficiency that still deletes the entire *Ddr* coding region (Figure 3D). We found that animals lacking any functional Ddr had impaired wrapping (Figure 3I-J, 3M; control: 79% normal morphology; *Ddr/Df*: 21% normal morphology; see Supplemental Figure S5 for examples of categorical scoring). Introduction of a single copy of a BAC clone containing the entire *Ddr* locus and a ∼17.4kb upstream region into the mutant strain restored normal wrapping morphology (Figure 3K, 3M; BAC rescue: 75% normal morphology), as did resupplying Ddr in wrapping glia via a *1xUAS-Ddr* construct driven by *nrv2-Gal4* (Figure 3L, 3M; *1xUAS* rescue: 70% normal morphology). Higher level overexpression using a *5xUAS-Ddr* construct caused morphological defects in a control background, which were not observed with the *1xUAS-Ddr* construct, suggesting that Ddr levels may need to be tightly regulated to promote normal wrapping and morphological development (Supplemental Figure S6). Together these results confirmed that Ddr is required cell autonomously in wrapping glia to regulate axon ensheathment.

We next analyzed nerves from control and *Ddr* mutant animals using transmission electron microscopy (TEM) and used the “wrapping index” (WI) metric to quantify ensheathment defects (Matzat et al. 2015). WI is equal to the number of individually wrapped axons plus the number of small bundles of axons, divided by the total number of axonal profiles and is expressed as a percentage. In control condition nerves, wrapping glia membrane separated axons in small bundles or individually (Figure 4A-B). The average WI in control conditions was ∼21% (Figure 4F: *nrv2* driver control: mean= 22.5%, n=26 nerves from 5 larvae, *w^1118^* control: mean= 21%, n=15 nerves from 3 larvae), consistent with previous reports (Matzat et al. 2015). We observed less separation of axons by wrapping glia membrane in *Ddr* mutants (Figure 4C-D): both *Ddr^41-2F^* homozygous and *Ddr^41-2F^*/*Df(2L)BSC186* animals exhibited a significant decrease in wrapping index (Figure 4F: *Ddr/Ddr* mean=14.8%, n=30 nerves from 4 larvae, p=0.0006 compared to *nrv2* control; *Ddr/Df(2L)BSC186*: mean= 11.9%, n=33 nerves from 4 larvae, p<0.0001 compared to *nrv2* control, one-way ANOVA with Dunnett’s multiple comparisons). These data confirm that loss of Ddr impairs axonal wrapping during development.

**Figure 4:**
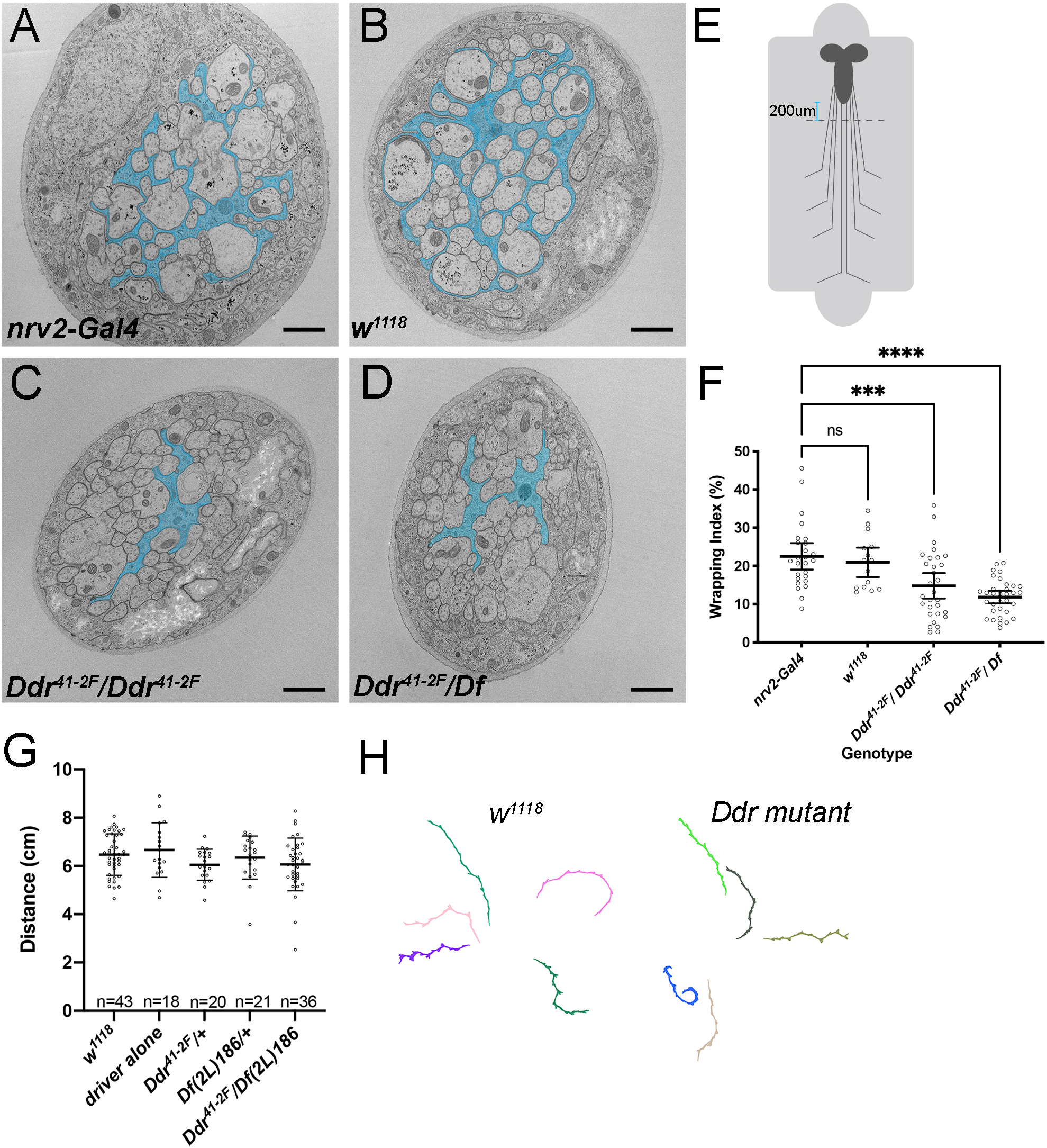
Loss of *Ddr* impairs axon ensheathment in larval nerves. (A) TEM cross section of an abdominal nerve from a control *nrv2-Gal4, UAS-CD8:GFP/+* 3^rd^ instar larvae. Wrapping glia membrane is pseudocolored cyan. (B) TEM cross section of an abdominal nerve from a control *w*^ll8^ 3^rd^ instar larva. (C) TEM cross section of an abdominal nerve from a *Ddr^41 - 2F^* homozygous mutant animal *(Ddr^41 - 2F^/Ddr^41 - 2F^ ; nrv2-Ga/4, UAS-CD8:GFP/+).* (D) TEM cross section of an abdominal nerve from a *Ddr* loss of function animal *(Ddr^41 - 2F^/Df(2L)BSC186; nrv2-Gal, UASCD8: GFP/+).* (E) Schematic oflarval fillets prepped for TEM. Sections for analysis are collected ∼200um from the distal tip ofthe VNC. (F) Quantification using the wrapping index (WI) metric. Wrapping index was reduced in Ddr loss of function conditions compared to control conditions in larval nerves. *Nrv2-Gal4, UAS-mCD8:GFP/+* controls (WI: 22.5%, n= 26 nerves, 5 larvae); *W ^1118^* wildtype background strain (WI: 21 %, n=15 nerves, 3 larvae, p=0.9546); *Ddr^-I-^* (WI: 14.8%, n= 30 nerves, 4 larvae, p=0.0006); *Ddr/Df* (WI: 11.9%,n=33 nerves, 4 larvae, p<0.0001); One-way AN OVA with Dunnett’s multiple comparisons against *Nrv2-Gal4.* Error bars: 95% Confidence interval. Scale bars = 1/-tm (G) Larval crawling, as measured by distance travelled per minute is not impaired in *Ddr* mutant animals compared to control conditions. One-way ANOVA with Tukey’s multiple comparisons, no significant differences between any conditions. n = # of larvae/condition. (H) Representative crawling paths from control and *Ddr/Df* mutant larvae.

To determine whether loss of Ddr specifically affected wrapping glia or more grossly disrupted the morphology of the nerve, we also assessed the other glial and axonal populations within the nerves of Ddr mutants. We did not observe any changes in either subperineurial or perineurial glia in TEM sections in the mutant conditions, nor differences in the total number of axon profiles observed (Supplemental Figure S7), suggesting that overall nerve assembly was relatively normal. Therefore, loss of Ddr seems to specifically alter wrapping without disrupting nerve morphology.

### Incomplete ensheathment of axons is not sufficient to alter crawling behaviors

We next sought to determine whether the impaired ensheathment in Ddr larvae was sufficient to impair behavior. We assayed larval crawling behavior in *Ddr* mutant third instar larvae again using the FIMTracker system (Risse et al. 2017). Comparing overall distance travelled over a one-minute period, we did not see any significant effects on basic crawling behavior between control and *Ddr* mutant larvae (Figure 4F-G), in contrast to the strong effects seen when wrapping glia were ablated. These results suggest that even the incomplete wrapping observed in *Ddr* mutants is sufficient to support basic neuronal functions during the ∼5 day period of larval life.

### Ensheathment of axons in an adult nerve is not impaired in *Ddr* mutants

Unlike ablation of wrapping glia, incomplete wrapping caused by the loss of Ddr did not significantly affect behavior or neuronal survival during the short larval period (∼5 days). However, since neuronal defects that are secondary to glial dysfunction can be age- or stress-related in vertebrate animals (Beirowski et al. 2014; Saab et al. 2016; Lappe-Siefke et al. 2003; Yin et al. 1998; Zöller et al. 2008), we investigated more deeply whether Ddr and proper wrapping were required for longer term support of axon health and function.

We turned our attention to the L1 sensory nerve in the adult wing. Sensory bristles and campaniform sensilla along the wing’s edge are innervated by peripheral sensory neurons located within the L1 wing vein. The axons of these neurons form the L1 sensory nerve that projects into the thorax (Figure 5A). The organization of the nerve is reported to be similar to larval nerves, with axons ensheathed by wrapping glia (Neukomm et al. 2014), but it had not previously been examined by TEM. We developed a protocol to perform TEM on the adult wing in order to examine the fine details of L1 wing nerve morphology. By imaging in the region distal to the fusing of the L3 nerve but proximal to the first sensory cell body along the anterior wing margin, we could reliably visualize all of the axons present in the L1 nerve and surrounding glia (Figure 5A, dashed red box). In control nerves at 5 **d**ays **p**ost **e**closion (dpe), we were surprised to find that all axons appear to be individually ensheathed by glial membranes and separated from one another (Figure 5B), instead of persisting in tightly compacted small bundles as in the larval nerves (compare to Figure 1A). When we examined *Ddr* mutant wings of the same age, we found no obvious defects in wrapping and axon separation between control and *Ddr* mutant wing nerves (Figure 5C-D), in contrast to the impaired wrapping we observed in *Ddr* mutant larval nerves. These data show that wrapping in the adult is more extensive than in larval nerves, and that ensheathment is completed during pupal stages or shortly after eclosion. Despite being more extensive and complete, wrapping in adult nerves does not strictly require Ddr. Ddr may be dispensable for wrapping in the adult, work redundantly with other molecules, or cause only a transient delay in wrapping that is rectified within a few days of eclosion.

**Figure 5:**
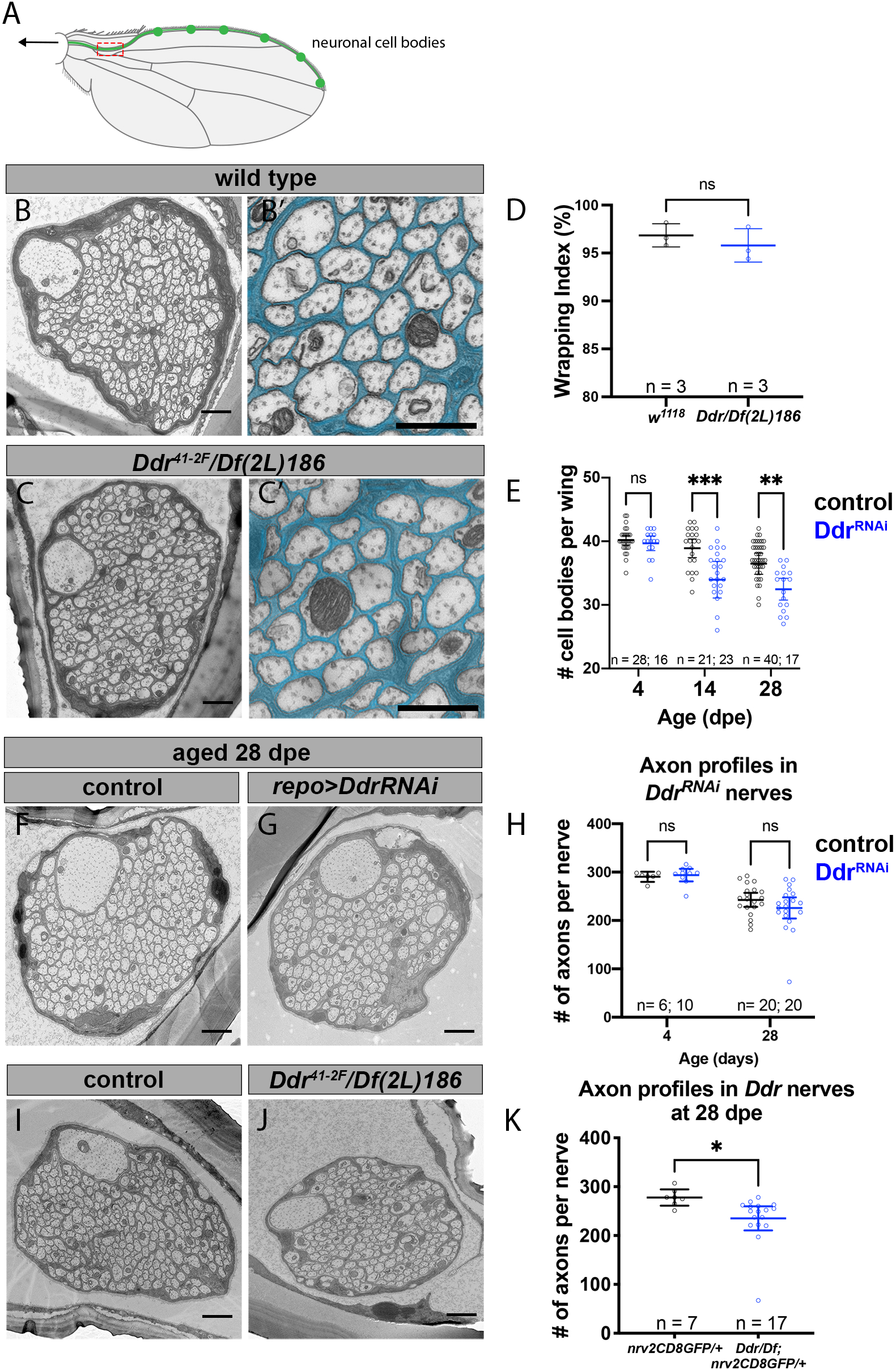
Loss of Ddr impairs long-term neuronal survival without affecting wrapping in adult nerves. **(A)** Schematic of the adult fly wing. The L1 vein lies along the anterior margin of the wing and contains ∼280 sensory neuron cell bodies, a subset are depicted in green. The axons of these neurons coalesce to form the L1 sensory nerve which projects into the body towards the fly CNS. The TEM imaging window is shown in red. In this location axons from all of the L1 sensory neurons are present, allowing for quantifications. **(B)** Cross section TEM of the L1 nerve shows that axons are separated from each other by glial membranes by 5 dpe. (B’) Magnified view from B with wrapping glia pseudocolored cyan. **(C)** Nerve morphology including wrapping is comparable in *Ddr* mutant nerves at this age, with axons separated and no obvious signs of impaired wrapping glia morphology. (C’) Magnified view from C with wrapping glia pseudocolored cyan. **(D)** Wrapping index of *Ddr* loss of function animals is unchanged compared to control. (Unpaired t-test, p=0.4378) **(E)** There is a reduction in the number of healthy glutamatergic GFP+ cell bodies in wings from aged *Ddr*^RNAi^ knockdown animals compared to age-matched controls. (Two-way ANOVA with Sidak’s multiple comparisons. 4dpe p=0.9774; 14dpe p=0.0007; 28dpe p=0.005) **(F)** Representative TEM image of a control nerve at 28 dpe shows normal wrapping/axon separation. **(G)** Representative *Ddr^RNAi^* knockdown nerve at 28 dpe also shows normal wrapping/axon separation. A wrapping glia nucleus is present in this section (asterisk). **(H)** Quantification of axon profiles in 28 dpe control and *Ddr* mutant animals shows a modest but significant reduction in axons in *Ddr* as compared to controls. (Unpaired t-test, p=0.0343) **(I)** Representative TEM of a control nerve from a 28 dpe animal shows normal wrapping. **(J)** Representative TEM of a nerve from a 28 dpe *Ddr* mutant animal also shows normal wrapping. **(K)** Quantification of axon profile number in control and *Ddr^RNAi^* glial knockdown nerves show no statistically significant difference in age dependent axon loss in contrast to our findings focusing on VGlut+ neuronal cell bodies by fluorescence (E). (Two-way ANOVA with Sidak’s multiple comparisons; 4 dpe p=0.9755; 28 dpe p=0.2519) Scale bars: B, C, F, G, I , J =1µm; B’& C’ = 600nm

### Loss of Ddr impairs the long-term survival of adult sensory neurons

Based on our observation of apparent neuronal loss in wrapping glia-ablated larvae, we speculated that long-term neuronal health in the adult L1 wing nerve might be impacted by loss of Ddr. To test this, we labeled the ∼40 glutamatergic neurons in the wing nerve using the *QF2/QUAS* binary system (*VGlut-QF2; QUAS-6xGFP*)(Potter et al. 2010), while at the same time using *Repo-Gal4* to knockdown *Ddr* in glia using RNAi. We examined wings at 4, 14, or 28 dpe and counted healthy GFP+ neuronal cell bodies along the wing margin at these timepoints from control and *Ddr-RNAi* animals to determine if loss of Ddr had any impact on long term neuronal survival. The number of neuronal cell bodies at 4 dpe was the same between control and *Ddr* knockdown animals suggesting that knockdown of *Ddr* did not affect neurogenesis (Figure 5E). Despite beginning with the same number of neurons (Figure 5E: 4 dpe mean cell body count = 40.2 vs. 39.7, p=0.9774), we found that there were significantly fewer healthy cell bodies in aged *Ddr*-*RNAi* wings compared to control at both ages examined (Figure 5E; 14 dpe mean cell body count 38.9 vs 34.0, p=0.0007; 28 dpe mean cell body count 36.5 vs 32.5, p=0.005, 2-way ANOVA with Sidak’s multiple comparisons). These results indicate that despite not overtly disrupting glial ensheathment, knockdown of *Ddr* in glial cells negatively impacts long-term neuron health and survival.

To confirm and extend these findings we used TEM to visualize L1 nerves from aged control and glial *Ddr-RNAi* animals. We collected cross sections from a region in which all axons should be present so we could compare the total number of axon profiles within the nerve. To mirror our fluorescent experiments, we used the same genotypes and we first examined wings at 4 dpe. The number of axon profiles in control nerves at 4 dpe showed little variation and was indistinguishable from *Ddr-RNAi* (Figure 5H: mean axon profile count 290.3±10 vs 293.9±18.3, p=0.98). In aged (28 dpe) control and *Ddr* knockdown wing nerves, although wrapping again appeared normal, there was overall more variability in axon profile counts (Figure 5F-H: control= 242.7±31.02, *Ddr-RNAi* =226±46.5 axon profiles), but there was no significant difference in axon profile count in the *Ddr-RNAi* condition (Figure 5E-G; p=0.25, 2-way ANOVA with Sidak’s multiple comparisons. This is in contrast to our findings in the same genotype when we focused exclusively on *VGlut-QF2*+ cell bodies (which marks ∼40 cell bodies out of ∼290 cell bodies total in the L1 nerve). We repeated these experiments using *Ddr* mutant animals and found a small but significant difference in the number of axon profiles (∼15% fewer axons) consistent with the size of our original findings with fluorescent labeling of a subpopulation of these neurons (Figure 5I-K; control =277.9 ± 17.9, *Ddr* = 235.2 ± 48.2, p=0.022, unpaired t-test). Since the effect was stronger in whole animal mutants, this may reflect insufficient RNAi knockdown, but we cannot rule out that loss of Ddr from neurons, or another tissue might contribute to this effect, or that only specific subsets of neurons are particularly vulnerable to Ddr loss. Nevertheless, these data indicate that glial Ddr modulates the long-term health of sensory neurons.

### Glial Ddr promotes increased axon caliber

Myelination can induce many changes in underlying axons including the redistribution of axonal proteins and increased axon caliber (Stassart et al. 2018), but whether non-myelinating ensheathment has similar effects is not known. One striking feature of the ultrastructure of the L1 wing nerve was the presence of a single, prominent, large caliber axon (Figure 5B). This axon is thus immediately identifiable in every sample, allowing us to directly compare its features across animals and conditions. A survey of the literature revealed that this axon likely belongs to the **d**istal **t**win **s**ensilla of the **m**argin (dTSM) neuron that innervates a campaniform sensillum (Dickinson and Palka 1987). A striking but unexpected observation was that the dTSM axon in *Ddr* mutant and knockdown conditions was significantly smaller than in controls even though it was still clearly identifiable as the largest axon in the nerve. We quantified axon caliber by measuring the cross-sectional surface area of this uniquely identifiable axon and comparing appropriate controls to *Ddr* mutant or glial *Ddr-RNAi* conditions from our 28 day aged wing TEM images. We found that in both glial-specific *Ddr*-knockdown and *Ddr* whole animal mutant nerves the caliber of the dTSM axon was significantly reduced compared to controls, revealing a role for glial Ddr in modulating axonal caliber (Figure 6A-F).

**Figure 6:**
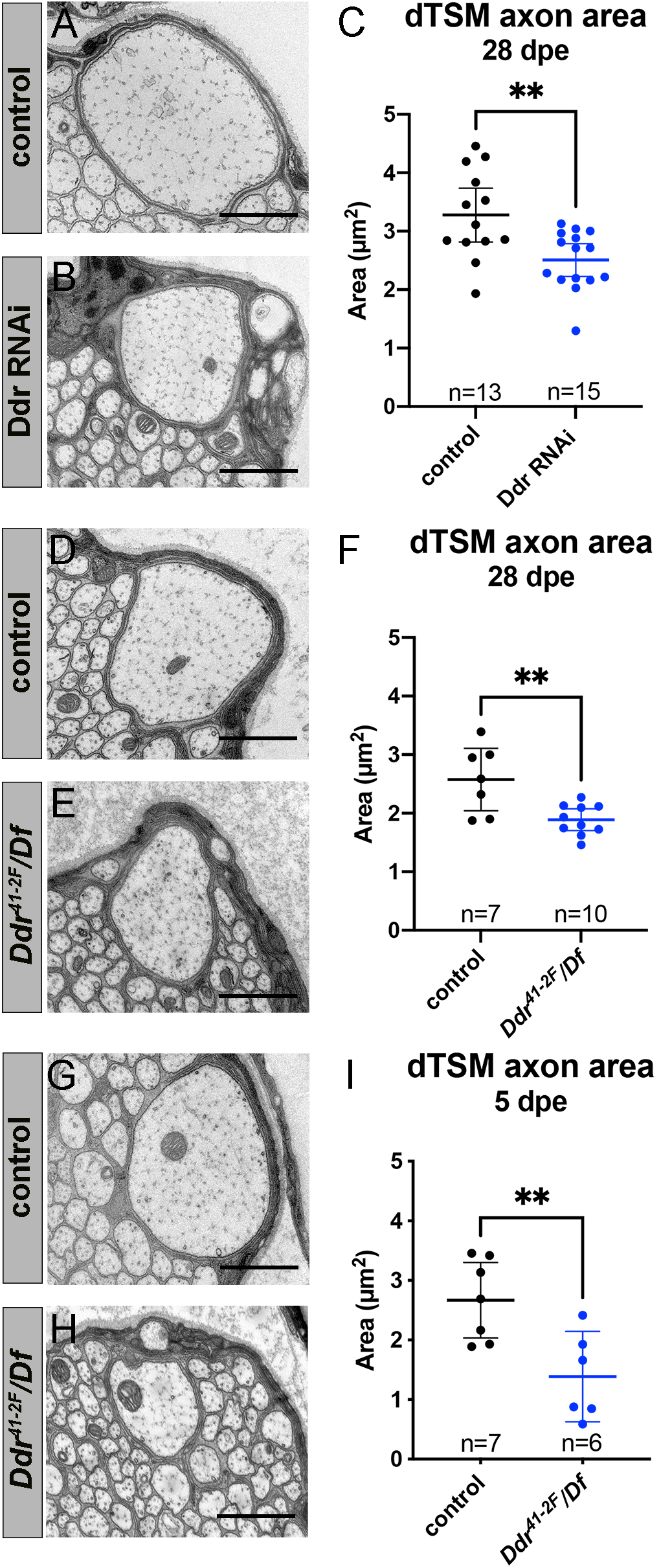
Ddr is required for normal axon caliber growth. (**A-C**) Axon caliber of the dTSM axons as measured by the cross-sectional area in TEM images is reduced in 28 dpe *Ddr^RNAi^*nerves compared to age-matched controls. (Unpaired t-test; p=0.0037) (**D-F**) Axon caliber of the dTSM axons as measured by the cross sectional area in TEM images is reduced in 28 dpe *Ddr* loss of function nerves compared to age-matched controls. (Unpaired t-test; p=0.0044) (**G-I**) Axon caliber of the dTSM axons as measured by the cross sectional area in TEM images is already reduced in 5 dpe *Ddr* loss of function nerves compared to age-matched controls suggesting that initial axon maturation is perturbed when Ddr is absent (Unpaired t-test; p=0.0073) Scale bars l µm.

To determine whether reduced caliber is due to reduced growth versus shrinkage, we examined dTSM caliber in control and *Ddr* mutant wings from 5 dpe and found that the reduction in size is even greater at this age (Figure 6G-I). dTSM axon caliber is reduced by ∼48% in *Ddr* mutants at 5 dpe (mean area: control = 2.670um^2^ and *Ddr* = 1.387um^2^) and ∼27% at 28 dpe (mean area control = 2.575um^2^ and *Ddr* = 1.890um^2^). We have examined a small number of L1 nerves from wild type freshly eclosed flies (<24hours post eclosion) and observed that all axons appear smaller than those from our standard 5 dpe timepoint suggesting that the first few days post eclosion are an important period of axon growth and maturation (Supplemental Figure S8). Together these data suggest that the reduced caliber we see in *Ddr* mutants is due to impaired growth, rather than shrinkage. Since knockdown of *Ddr* selectively in glia is sufficient to cause a similar reduction in caliber to what is observed in whole animal mutants (∼23% reduction at 28 dpe in RNAi experiments) our data identifies a cell non-autonomous role for glial Ddr in the control of axon caliber.

### *Ddr* interacts with *Mp* to promote wrapping in larval nerves

We next sought to understand the molecular mechanism(s) by which Ddr is activated to promote axon ensheathment. Mammalian Ddrs are activated by collagens *in vitro* and are thus considered non-canonical collagen receptors (Vogel et al. 1997). We wondered if *Drosophila* Ddr also interacts with collagen(s) to mediate its effects on wrapping in the larvae. The *Drosophila* genome contains 3 collagen genes. One of them, *Multiplexin* (*Mp*), was identified as a potential hit that affected wrapping glia morphology in third instar nerves in our initial screen. Knockdown of *Mp* using *nrv2- Gal4* caused moderate to severe wrapping defects when analyzed by confocal microscopy (Figure 7A-B). We studied expression of Mp protein in wild type nerves by using a MiMiC-GFP line in which the endogenous Mp protein is tagged with GFP. We observed punctate GFP expression throughout larval nerves suggesting that Mp, a secreted protein, is expressed by one or more of the cell types within the nerve (Figure 7C).

**Figure 7:**
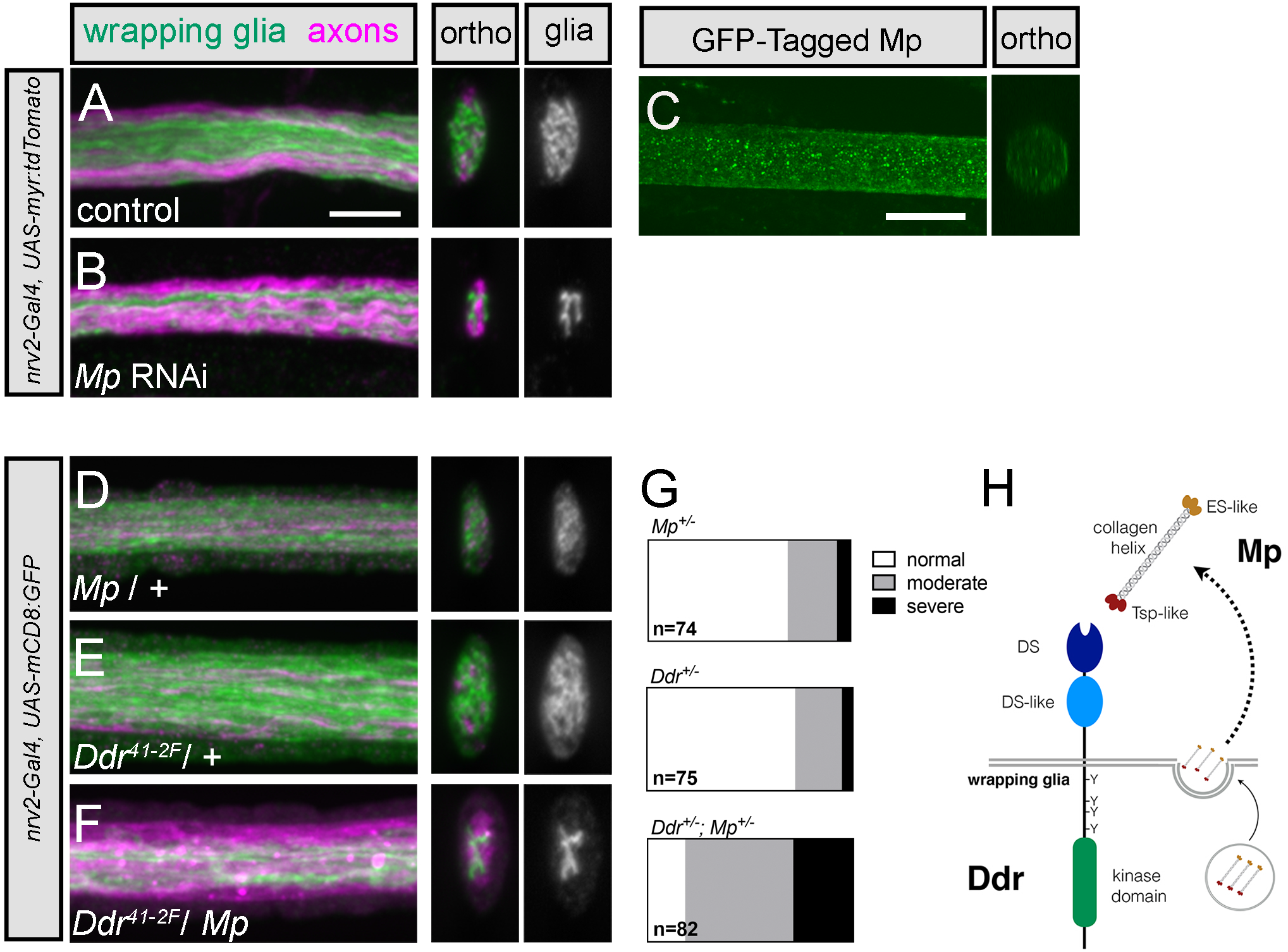
*Mp* genetically interacts with *Ddr* to promote wrapping in larval nerves. (**A-B**) Representative images of control and *Mp* RNAi knockdown in larval nerves. *Nrv2-Ga14* driven tdTomato is psuedocolored green. A subset ofaxons is labeled with anti-futsch antibody (magenta). Compared to control nerves, RNAi conditions show large areas of the nerve cross section without glia membrane coverage. (**C**) Mp-GFP expression in larval nerves. A MiMiC construct that adds a GFP tag to Mp protein is localized throughout larval nerves. (**D-F**) *Mp* and *Ddr* heterozygous nerves show normal wrapping glia morphology with good glial coverage of the nerve cross section indicating that one copy of each gene is sufficient to promote wrapping. In double heterozygous animals missing just one copy of each gene, however, coverage ofthe cross section is reduced indicating that wrapping is impaired. (**G**) Categorical scoring of *Ddr* and *Mp* single and double heterozygotes. n= # of nerves analyzed from 9-11 larvae per condition. (**H**) Proposed model of Mp and Ddr interaction. Based on the strong phenotype observed when Mp is knocked down specifically in wrapping glia, we hypothesize that wrapping glia secrete Mp which can act as autocrine activator of Ddr to drive normal wrapping glia morphogenesis.

To determine whether *Ddr* and *Mp* might be working in the same genetic pathway to modulate wrapping in larval nerves, we crossed heterozygous *Ddr* and *Mp* mutants together and analyzed the double heterozygous progeny for defects in larval wrapping glia morphology. While animals that were heterozygous for either gene on its own did not show defects in wrapping glia morphology, we found that the double heterozygous animals exhibited moderate to severe defects in glial morphology (*Ddr/+*: 77% normal morphology; *Mp/+:* 72% normal morphology; *Ddr/Mp*: 18% normal morphology; Figure 7D-G). These data suggest a strong genetic interaction between the *Ddr* and *Mp* loci and imply they might function in a common genetic pathway to control wrapping glia morphological development. Given that RNAi knockdown of *Mp* specifically in wrapping glia is sufficient to cause a wrapping defect, our data suggest a model in which Mp acts in an autocrine fashion to activate Ddr and drive downstream signaling to promote wrapping (Figure 7H).

## Discussion

Non-myelinating ensheathment of axons is a conserved but understudied feature of peripheral nervous systems. Although this type of multi-axonal ensheathment has been less studied as compared to myelination, a growing body of evidence indicates it is important for the health and function of neurons and axons in the periphery. For example, Schwann cell specific loss of the transmembrane receptor LDL receptor related protein-1 (LRP1) causes both thin myelin and abnormal Remak bundle structure. These cKO animals also showed a lowered pain threshold, suggesting that the physiology of nociceptor neurons is impaired when Remak ensheathment is disrupted (Orita et al. 2013). Disrupting metabolism in SCs causes progressive axon loss, with small unmyelinated fibers dying first, before myelinated fibers begin to show signs of degeneration (Viader et al. 2013, 2011; Beirowski et al. 2014). In the fly, disruption of axonal wrapping leads to uncoordinated behavioral responses that hint at aberrant ephaptic coupling between neighboring axons in nerves when not properly separated (Kottmeier et al. 2020). Such coupling could cause inappropriate activation of sensory or nociceptive neurons underlying peripheral neuropathies.

Previous studies from our lab have shown that wrapping glia are required to clear neuronal debris after nerve injury and mediate injury signaling between injured and intact “bystander” neurons which might be important for functional recovery after nerve trauma (Neukomm et al. 2014; Hsu et al. 2021). These and other findings suggest that Remak-type ensheathment and axon-glia signaling of unmyelinated fibers plays a variety of underappreciated roles in peripheral nerve physiology that contribute to the pathophysiology of a number of PNS disorders including debilitating peripheral neuropathies and responses to nerve injury.

In order to gain insight into non-myelinating ensheathment, we used the *Drosophila* peripheral nerves to identify a novel molecular pathway important for the development and function multi-axonal ensheathment. We generated a new Split-Gal4 intersectional driver to specifically target wrapping glia for functional and behavioral studies to better understand how wrapping glia support axon health, physiology and ultimately circuit function. Finally, we uncovered roles for glia in mediating long-term neuronal survival and driving increased axon caliber that are separable from overt effects on wrapping, demonstrating that non-myelinating ensheathing glia perform critical, previously unappreciated, roles in nervous system development, maintenance, and function.

### Ddr and Mp regulate wrapping in larval nerves

A main advantage of *Drosophila* is the ability to conduct large scale *in vivo* screens. We made use of available *UAS-RNAi* libraries to broadly screen for novel regulators of axonal ensheathment in intact nerves. Our morphological screen was sensitive enough to identify genes previously implicated in wrapping glia development including *vn*, *LamB1*, and *mys*, validating our approach. Moreover, in the case of *Ddr*, we were able to identify an important novel regulator of ensheathment that a simple behavioral or lethality screen would have missed in light of our follow-up behavioral testing. Knockdown of *Ddr* specifically in wrapping glia resulted in reduced glial membrane coverage in nerve cross sections by fluorescence microscopy. Similar phenotypes were observed in *Ddr* loss-of-function animals and could be rescued by resupplying Ddr specifically in wrapping glia, confirming the specificity of our RNAi results. TEM clearly showed that reduced glial membrane coverage at the light level corresponds to decreased axon wrapping. Unexpectedly, we observed that strong overexpression of *Ddr* using a *5XUAS-Ddr* construct in a control background caused frequent morphological defects in wrapping glia that were not observed when using a *1XUAS-Ddr* construct, suggesting that Ddr signaling must be tightly regulated to successfully promote wrapping.

Though neither of the vertebrate homologs, Ddr1 or Ddr2 has been explicitly implicated in glial development, several lines of evidence suggests that Ddr1 may have a conserved role in vertebrate glial development or function. Ddr1 is highly expressed in the mouse OL lineage starting from when the cells begin to associate with axons, is upregulated in newly formed OLs after cuprizone treatment, and is expressed in both myelinating and Remak Schwann cells (Gerber et al. 2021; Zhang et al. 2014; Franco-Pons et al. 2006, 2009). Moreover, Ddr1 is expressed in human oligodendrocytes and myelin, and variants in the human gene have been correlated with abnormal white matter and schizophrenia (Roig et al. 2010; Gas et al. 2018).

Vertebrate Ddr1 and Ddr2 are potently activated by collagens in vitro (Vogel et al. 1997), prompting us to investigate whether collagens were involved with Ddr function in fly nerves. We had discovered that loss of the *Drosophila* collagen Mp disrupted glial morphology in our initial RNAi screening, and so investigated whether these genes might work together to promote ensheathment. Together with the established roles for Ddr1 and Ddr2 as collagen receptors, the strong genetic interaction we observed between *Ddr* and *Mp* is consistent with a model in which Mp acts as a collagen ligand for Ddr during axonal ensheathment. Our expression data from the *Mp-GFP* protein trap showed expression throughout the nerve. Although previous reports indicate that Mp is also expressed in one or both of the outer peripheral glia layers that extend beyond the nerve to cover NMJ synapses (Wang et al. 2019), the strong ensheathment defect seen when *Mp* is knocked down exclusively in wrapping glia indicates that wrapping glia themselves are the primary source of the Mp required for wrapping glia morphogenesis.

We hypothesize that Mp can act an autocrine collagen ligand to activate glial Ddr to drive wrapping in larval nerves (Figure 7H). This parallels developing Schwann cells, which secrete their own basal lamina, several components of which acts as regulators of their development including laminins and collagen. For example, laminin-211 serves as a ligand for GPR126 to promote myelination (Petersen et al. 2015). Mp is the sole *Drosophila* homolog of collagen types XV/XVIII, containing a central helical collagen region with a cleavable N-terminus thrombospondin like domain and C-terminal endostatin like domain (Meyer and Moussian 2009; Myllyharju and Kivirikko 2004). Collagen 15a1 and 18a1 are expressed in mouse peripheral nerves and *col15a1* mutants have radial sorting defects, suggesting that *Mp*’s role in promoting axon wrapping is likely conserved (Rasi et al. 2010; Gerber et al. 2021; Chen et al. 2015). In fact, *Mp* appears to play multiple important roles in nerve biology. Mp secreted by the outer glia layers acts via its cleaved endostatin domain, which is essential for homeostatic plasticity at motor neuron synapses (Wang et al. 2014, 2019). How Ddr activation within wrapping glia ultimately drives axon wrapping still remains to be determined, but Ddr joins two other RTKs—EGFR and FGFR—as important and conserved regulators of axon ensheathment (Matzat et al. 2015; Kottmeier et al. 2020; Franzdóttir et al. 2009).

### Wrapping glia are required for normal larval behavior

The *nrv2-Gal4* driver has been the standard method to genetically target wrapping glia but it is imperfect for manipulation of wrapping glia in ablation or behavioral assays due to its expression in several subtypes of CNS glia. We generated a new Split-Gal4 intersectional driver that drives exclusively in wrapping glia. This allowed us to perform more precise wrapping glia ablation. We found that genetic ablation of wrapping glia led to severely impaired larval locomotion, indicating the wrapping glia are essential for basic crawling circuit function. This phenotype was particularly striking in light of that fact that we did not observe any clear crawling defect in *Ddr* mutant larvae, even though wrapping was severely impaired. There are several possible explanations for these observations. First, it may be possible that non-contact mediated mechanisms, such as one or more secreted factors constitute wrapping glia’s essential contribution to axon health and physiology.

Alternatively, perhaps even a small amount of direct glia-axon contact at any point along the axon length may be sufficient to support axon function. This would be consistent with the lack of overt behavioral defects in newly-hatched 1^st^ instar larvae, which have poor wrapping compared to later stages (Matzat et al. 2015; Stork et al. 2008), and even in wild type third instar larvae, in which not every axon is individually wrapped. This supports the notion that at least in the larvae, wrapping per se may not be strictly required for very simple behaviors, but glial presence and/or metabolic support and perhaps at least some contact is sufficient to support function for the ∼5 days of larval life. Our results are similar to what has been recently reported (Kottmeier et al. 2020) using a different approach to target wrapping glia for ablation, where only minor behavioral defects were observed when disrupting FGFR signaling but profound crawling defects were seen upon ablation.

Together these data support the conclusion that even limited wrapping or simply some degree of glia-axon contact is sufficient to support axon survival and nerve function compared to no glia at all in larval nerves.

### Loss of Ddr impairs long-term neuronal survival

Previous studies of oligodendrocytes and Schwann cells have found that impairing glial function can result in seemingly normal wrapping and circuit function in young animals, with deficits only appearing when the system is stressed or aged (Saab et al. 2016; Beirowski et al. 2014; Lappe-Siefke et al. 2003; Zöller et al. 2008). Studying wrapping in adult *<u>Drosophila</u>* allows for aging and maintenance studies that the short larval period precludes. Adult peripheral nerves are encased in a transparent but hard cuticle that allows for live-imaging but makes fixation challenging. Because of the resolution limits of light microscopy, we established a reliable method to study their ultrastructure using TEM. We found that the wrapping of axons by glia in the adult wing nerve is strikingly different from the larva, as all axons appear to be separated and individually ensheathed by glial membranes. To our surprise, wrapping was not obviously impaired in adult nerves of *Ddr* knockdown or mutant animals. Whether this is because it causes only a transient delay during pupal development, or is simply not required for adult wrapping glia morphogenesis remains unclear. In larval wrapping glia, there are 3 RTKs (EGFR, the FGFR heartless, and now Ddr) that are each required for normal ensheathment, and thus cannot fully compensate for one another (Matzat et al. 2015; Kottmeier et al. 2020). One difference between larval and adult wrapping glia is the size of each cell. One larval wrapping glia cell covers the majority of the nerve length from where it exits the VNC to the territory of the second wrapping glia which has its cell body where nerve enters the muscle field. This means that a single wrapping glial cell must undergo tremendous growth to both keep up with longitudinal growth as the animal grows and elongates the nerve, as well as radial growth to separate axons. A single cell can end up covering from ∼750um-2.5mm of nerve length depending on the segment. In the wing nerve, there are ∼15 wrapping glia along the region of the L1 nerve we analyze, which is ∼400um long. Perhaps in the larval system the cell is pushed to its growth limits and any perturbation in pro-wrapping signaling has a strong effect of morphology, whereas in the adult nerve the system is robust and redundant enough to withstand perturbations of single genes.

Loss of *Ddr* led to spontaneous neurodegeneration in the nerve as animals naturally aged. Such an uncoupling of neuron health from overt effects on myelination has been demonstrated previously. For example, *Cnp1* mutant mice show severe age-dependent neurodegeneration although they have grossly normal myelin with only subtle changes in myelin ultrastructure (Lappe-Siefke et al. 2003; Snaidero et al. 2017), and loss of PLP results in axon degeneration despite having largely normal myelin (Garbern et al. 2002; Klugmann et al. 1997; Griffiths et al. 1998). We found that the number of VGlut+ neurons was reduced in aged wings of *Ddr* knock down animals, indicating that glial *Ddr* is important for long-term neuronal survival. When we analyzed *Ddr* whole animal mutants by TEM we found a small but significant reduction in axon profile number, which should correspond to the number of surviving neurons. The effects were modest, and together with the increased variability observed suggests that absence of Ddr signaling increases the susceptibility of subpopulations of neurons to insult or injury that may underlie age-related degeneration.

### Glial Ddr regulates axon caliber

Myelination can directly affect the structure and function of the axons they wrap, including controlling caliber. In general, myelination increases caliber. For example, dysmyelinated *Trembler* mice have reduced axon calibers compared to controls (Waegh et al. 1992) and in the PNS caliber along a single axon can vary with reduced caliber at points without direct myelin contact, such as nodes of Ranvier (Hsieh et al. 1994). Axon caliber is an important determinant of conduction velocity but varies widely between neuronal subtypes, so achieving and maintaining appropriate caliber for each axon during development is likely to be critical for proper circuit function. How non-myelinating ensheathment might impact axon caliber has not been as clearly established. Here, we find that glial Ddr promotes increased axon caliber of a non-myelinated axon. We focused on the dTSM axon, which can be reliably identified across animals, so we could directly compare caliber between conditions. The reductions we observed in axon caliber were similar between *Ddr* mutants and when Ddr was specifically lost from glia cells, supporting a non-cell autonomous role for glial Ddr in regulating axon caliber. The effect is considerable—nearly a 50% reduction in axon caliber at 5 dpe. We hypothesize that by this time point, wildtype dTSM axons have reached their mature caliber, as it is comparable between 5 dpe and 28 dpe in comparable genetic backgrounds. In *Ddr* mutants, however, we observe that the relative size compared to controls changes over time, suggesting that in *Ddr* mutants (or knockdowns) the axon continues to increase its caliber, perhaps in an effort to achieve the optimal size, though the axons still remain ∼25% smaller than wild type axons at 28 dpe.

Two proteins, MAG, which acts to increase caliber of myelinated axons (Yin et al. 1998), and CMTM6, which restricts caliber of myelinated and unmyelinated axons (Eichel et al. 2020) are the only proteins reported to cell non-autonomously affect caliber of vertebrate axons. In the fly, a shift in the average size of axons in larval nerves is observed when wrapping glia are absent or severely disrupted supporting a general role for wrapping in promoting axon size (Kottmeier et al. 2020). We show that Ddr is required for increased axon caliber even when wrapping appears intact.

The exact molecular mechanism by which Ddr may promote increased caliber size remains unclear. The control of axon caliber, generally, is not well understood, with few studies looking at either cell autonomous or non-autonomous regulators. Genes involved in the general regulation of cell size have been implicated as cell autonomous determinants. For example, in the fly, S6 kinase signaling is a positive regulator of motorneuron size, including axon caliber (Cheng et al. 2011). In mammalian axons, it is believed that the phosphorylation state of neurofilaments and microtubules ultimately determines their spacing, thereby contributing to control of caliber (Yin et al. 1998). It seems unlikely that *glial* Ddr can directly phosphorylate *axonal* microtubules, so determining how Ddr activity in glia influences the axonal cytoskeleton to control caliber growth is an important next step. A 25-50% reduction in caliber would be predicted to impact conduction velocity along the dTSM axon and given that campaniform sensilla provide essential rapid sensory feedback to fine tune movement, it will be of interest to test conduction velocity and/or flight behavior directly in *Ddr* mutant animals to see how the proprioceptive circuit might be affected.

Taken together, our studies identify Ddr as an important regulator of wrapping glia development and function in the fly, with distinct roles in larval and adult wrapping glia. Ddr is essential for the normal morphological development of axon wrapping in the larvae, and later it mediates important axon-glia communication that controls axon caliber growth and affects neuronal health and survival. Given its expression pattern in vertebrate oligodendrocytes and Schwann cells, it seems likely these essential functions are conserved in vertebrates. Further study into how Ddr functions in both fly and vertebrate glia promises to increase our understanding of axon ensheathment in health and disease.

## Methods

### Drosophila genetics

*Drosophila melanogaster* were raised according to standard laboratory conditions. Larval RNAi experiments, including controls, were conducted at 29°C to increase expression of the UAS-RNAi constructs. All other experimental crosses and aging experiments were conducted at 25°C. A full list of all fly strains used in this study can be found as a table in the Supplemental Methods. Fly strains generated for this study include *Ddr^41-2F^*, *Ddr^13-1M^, WG-SplitGal4, 5XUAS-Ddr* and *1XUAS- Ddr.* Full details on their construction are also in the Supplemental Methods.

### Immunohistochemistry and Confocal Analysis

For confocal fluorescent imaging, larvae were filleted on sylgard plates in room temperature (RT) PBS and fixed on a shaker in 4% paraformaldehyde (Electron Microscopy Sciences, EMS) in PBS for 20 minutes. After washing with PBS and PBST (0.3% Triton-X), larvae were incubated in primary antibodies overnight at 4°C with gentle agitation on a shaker. After washing in PBST at RT on a shaker, secondary antibodies were added and incubated overnight at 4°C on a shaker, shielded from light. After washes in PBST, samples were mounted on glass slides using Vectashield (Vector Labs). The following primary antibodies were used in this study: chicken *α*-GFP (1:1000, Abcam, ab13970); rabbit *α*-DsRed (1:600, Clontech, #632496); rat *α*-mCherry (1:2000, Invitrogen, #16D7); mouse *α*-futsch (22C10, 1:500, DSHB); mouse *α*-Repo (1:100, DSHB); goat *α*-HRP pre-conjugated to Alexa488, CY3, or CY5 (1:100, Jackson Immuno Research); rabbit *α*-Oaz (1:5000, this study, see details below). Primary antibodies were detected with the appropriate donkey secondary antibody conjugated to DyLight 488, Alexa488, Cy3, Rhodamine Red-X, CY5, or DyLight 405 reconstituted per the manufacturer’s instructions and use at final dilutions of 1:250 (Jackson Immuno Research). Phalloidin-iFluor 647 and 405 were used at 1:2500 (Abcam).

Confocal images were obtained on either an Innovative Imaging Innovations (3i) spinning-disk confocal microscope equipped with a Yokogawa CSX-X1 scan head and Hamamatsu camera with SlideBook software (3i) or on a Zeiss spinning disk confocal microscope equipped with a Yokogawa CSX-X1 scan head and Hamamatsu camera with Zen software. Images were obtained with 10x air, 40x 1.3NA Oil, or 63x 1.4NA Oil objectives. Confocal stacks taken to analyze wrapping glia morphology in nerve cross sections were taken with the 63x objective, using the recommended optimal slice size of 0.27 microns.

To score wrapping glia morphology, images were obtained from regions ∼200 microns from the tip of the VNC (to roughly correspond to the TEM analysis location). At 63x magnification ∼100um of nerve length could be analyzed in an image. Orthogonal sections were visualized using SlideBook or Zen software and the cross section of each nerve was examined over the length in the captured image. Morphology was scored categorically as being “normal,” having “moderate” defects in nerve coverage, or having “severe” defects in nerve coverage. Nerves were not scored if the Z axis of a particular nerve was too compressed to visualize morphology well. Representative orthogonal views were exported to prepare figures. See Figure 2—figure supplement 1 for images and descriptions of each category. The number of larvae and nerves scored for each quantified genotype is included in the text or appropriate Figure legend.

### Electron microscopy

Third instar larvae were manually processed for TEM as dissected fillets, using standard TEM processing procedures. Drosophila wings were processed using microwave-assisted fixation with a protocol adapted from (Czopka and Lyons 2011; Cunningham and Monk 2018) protocols for zebrafish larvae, with the main differences being the elimination of all 10 minute RT “hold” steps. After fixation and embedding in Embed-812, 70nm sections were collected on 200 mesh copper grids (larvae) or 100 mesh Formvar film coated grids (wings), counterstained with UA and lead citrate and imaged on a Tecnai T12 electron microscope operating at 80kV or 120kV accelerating voltage equipped with an AMT digital camera and software. Full details of fixation and processing are available in the Supplemental Methods.

### Aging assay and live imaging of wings

Animals of appropriate genotypes were crossed at 25°C. Correct progeny were selected using visible markers and aged in small groups in cornmeal agar vials for the indicated number of days at 25°C. Flies were transferred to fresh food vials every 3-7 days. On the appropriate day (4, 14, or 28 days after collection), flies were anesthetized and one wing was removed from each animal using fine dissection scissors (Fine Science Tools). Wings used for fluorescence imaging were mounted in Halocarbon oil 27 (Sigma) on glass slides for live imaging as described in (Neukomm et al. 2014). For the neuronal survival assay, wings were inspected for tears and injuries along the L1 vein at 63x with transmitted light and only wings with no visible physical damage were assessed further. GFP+ cell bodies were visualized and counted using a Zeiss Spinning Disk confocal system (Axio Examiner equipped with a Yokogawa spinning disk, Hamatsu camera, and Zen software) using a 63xNA1.4 oil objective. Cells were counted as intact if they had a clear nucleus and dendrite or were considered dead if they were shrunken and the dendrite or nucleus were not clearly visible. Wings to be analyzed by TEM were immediately placed in fixative and processed as described in the Supplemental Methods.

### Larval Tracking

Larval crawling behavior was analyzed using the frustrated total internal reflection imaging method and FIMTrack software (Risse et al. 2017). Briefly, larvae were washed briefly in ddH20 and kept on agar plates prior to testing. Approximately 5 larvae of the same genotype were placed near the center of a 0.8% agar surface positioned above an IR camera using a paintbrush. Behavior was recorded at RT at 10 frames/second for 1 minute. Videos were analyzed using FIMTrack software to extract data about individual larva. Crawling paths for each larva were automatically generated by FIMTrack and information of total distance travelled for each larva was exported to Excel and converted from pixels to cm before being analyzed using GraphPad PRISM software. Only larvae that were successfully tracked for the full minute were included in the analysis.

### Analysis and Statistics

TEM image analysis was performed in FIJI/ImageJ using an AMT plugin to import image metadata. Wrapping index (WI) was quantified as described in Matzat et al, 2015. Briefly, the Cell Counter plugin was used to manually tally the total number of axons and then manually tally the number of bundles or singly wrapped axons. WI was then computed for each nerve as: # of singly wrapped + # axon bundles/Total # of axons, expressed as a percentage. Axon counts in wing nerves were also performed manually with the Cell Counter plugin in FIJI. dTSM cross sectional area was performed using the polygon selection tool to manually trace the circumference of the dTSM axon and then using the Measure Tool to obtain the cross-sectional area that had been outlined. Nerves were excluded from an analysis if knife marks, resin folds, or debris blocked the feature(s) needing to be measured or if the section angle was too far from perpendicular resulting in elongated and blurry axonal profiles. For larval experiments, both male and female larvae were used unless the genetics of a specific experiment only allowed for one sex to be used (e.g. transgene on the X chromosome coming from a male parent). For adult experiments only female flies (wings) were used. Wing size and the number and type of sensory neruons present in the wing are sexually dimorphic, which necessitated using a single sex to compare axon number and caliber. Females were selected because their wings are slightly larger and thus easier to handle, and this facilitates genetic experiments that involve genes and/or transgenes on the X chromosome.

Statistical analysis was performed using GraphPad Prism software. The appropriate statistical test was determined based on the experimental design and how many conditions were being compared. Details on the exact sample sizes and statistical tests used are included in the corresponding Results section and/or Figure.

## Acknowledgements

We are grateful to Dr. Christian Klambt, Dr. Joshua Dubnau, the Bloomington Stock Center, and the Vienna Drosophila Resource Center for providing fly stocks. We would like to acknowledge FlyBase.org for being an invaluable research tool for *Drosophila* scientists (Larkin et al. 2020). Anti-futsch (22C10), anti-Repo (8D12), and anti-Elav (7E8A10) monoclonal antibodies developed by Drs. Seymor Benzer, Corey Goodman, & Gerald Rubin (respectively) were obtained from the Developmental Studies Hybridoma Bank created by the NICHD of the NIH and maintained at The University of Iowa, Department of Biology, Iowa City, IA 52242. We thank Dr. Michael Brodsky for assistance with CRISPR-Cas9 mediated mutagenesis. We thank Dr. Lara Strittmatter and the staff of the University of Massachusetts Medical Center Electron Microscopy Facility; Dr. Robert Kayton and Dr. Lisa Vecchiarelli of the OHSU Electron Microcopy Facility; and Dr. Deborah Hegarty for their expertise and assistance with TEM protocol development and equipment. We are indebted to Dr. Kelly Monk and the Monk lab for TEM advice and generous sharing of their microwave tissue processing equipment. We thank Rachel Bradshaw for her help with the RNAi screen and Alex Larson for her assistance with TEM prep work. We thank Drs. Graeme Davis and F. Javier Bernardo-Garcia for discussions and critical reading of the manuscript. Finally, we thank all members of the Freeman lab for their advice and feedback throughout this project. This work was supported by NINDS P30 NS061800 (SAA) and NINDS R01NS112215 & R01059991(MRF). MRF was an investigator with the Howard Hughes Medical Institute during a portion of this study.

## Author Contributions

M.M.C.: Conceptualization, Methodology, Formal Analysis, Investigation, Data Curation, Writing-Original Draft, Visualization, Project Administration, Funding Acquisition

A.P.L.: Methodology, Formal Analysis, Investigation, Data curation, Writing-Review and Editing, Visualization

J.Q.H.: Methodology, Investigation, Writing-Review and editing

A.E.S.: Investigation

S.A.A.: Resources, Writing-Review and Editing, Funding Acquisition

M.R.F .: Conceptualization, Resources, Generation of the anti-Oaz antibody, Writing-Review and Editing, Project Administration, Supervision, Funding Acquisition

