## Supplemental Figs & Methods for "*Discoidin domain receptor* regulates ensheathment, survival, and caliber of peripheral axons"

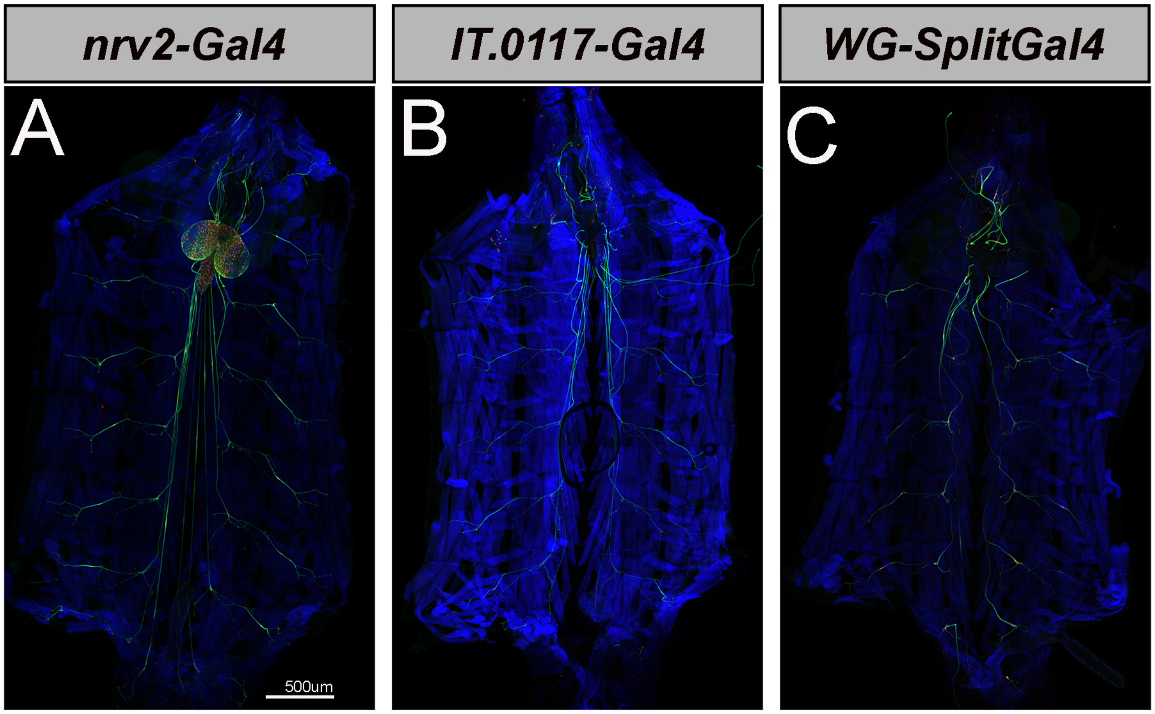

**Supplemental Figure S1: *WG-SplitGal4* whole body expression pattern**
**(A)** Whole animal expression pattern of *nrv2-Gal4* driving *UAS-CD8:GFP* (green) and *UAS-mCherry*$^{nls}$ (red). Body wall muscles are labelled with phalloidin (blue). **(B)** Whole animal expression pattern of *IT.0117-Gal4* driving *UAS-myrGFP* (green) and *UAS-H2B:mCherry* (red), **(C)** Whole animal expression pattern of *Wrapping glia-SplitGal4* (made by combining *nrv2-Gal4*$^{DBD}$ and *IT.0117-Gal4*$^{VP16AD}$) driving *UAS-mCD8:GFP* (green) and *UAS-mCherry*$^{nls}$ (red). *WG-SplitGal4* drives exclusively in wrapping glia in the periphery without any evidence of neuronal or glial expression in the CNS nor in any other tissue in the larva.

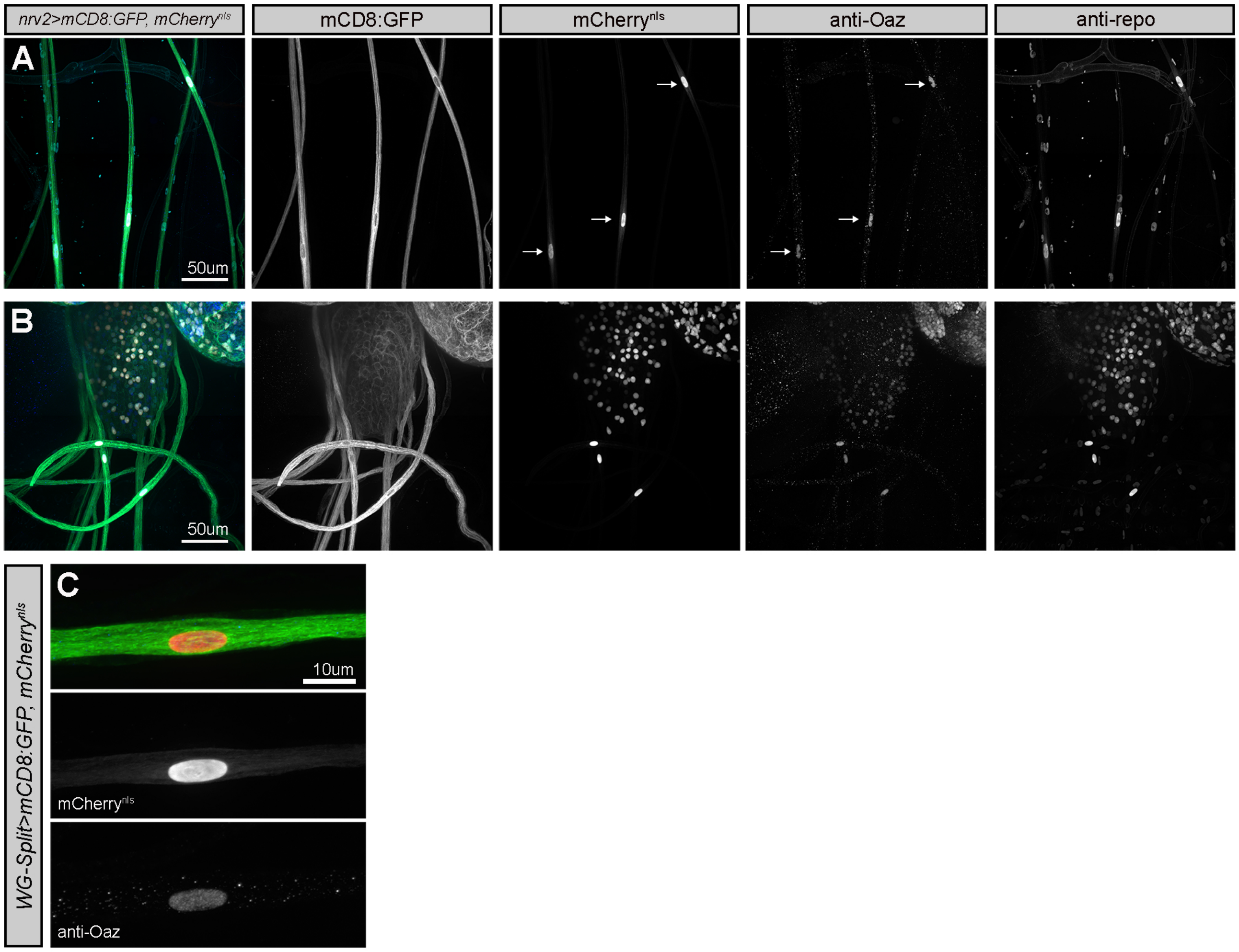

**Supplemental Figure S2: The transcription factor Oaz can be used to label wrapping glia nuclei**
**(A)** Confocal projection image of nerves from a *nrv2>mCD8:GFP, mCherry$^{nls}$* larva. First panel merged image showing: membrane GFP (green), nuclear mCherry (red), anti-Oaz (blue), anti-Repo (cyan). Channels are separated out in subsequent panels. *Nrv2>mCherry* positively identifies wrapping glia nuclei. Only these Cherry+ nuclei, and not other Repo+ nerve glia nuclei are positive for anti-Oaz staining (arrows). **(B)** Confocal projection image of the CNS and proximal nerves from a *nrv2>mCD8:GFP, mCherry$^{nls}$* larva. Channels are separated out as in A. Along the nerves, only wrapping glia nuclei are labelled with anti-Oaz. Oaz is also expressed in CNS nuclei, including a subset of the glia within the *Nrv2-Gal4* expression and also some neurons. Not shown, but faint staining is also observed in muscle nuclei. **(C)** Confocal projection of a single nerve from a *WG-SplitGal4>mCD8:GFP, mCherry$^{nls}$* larva showing co-localization of anti-Oaz with a mCherry positive nucleus, providing additional confirmation that *WG-SplitGal4* labels wrapping glia.

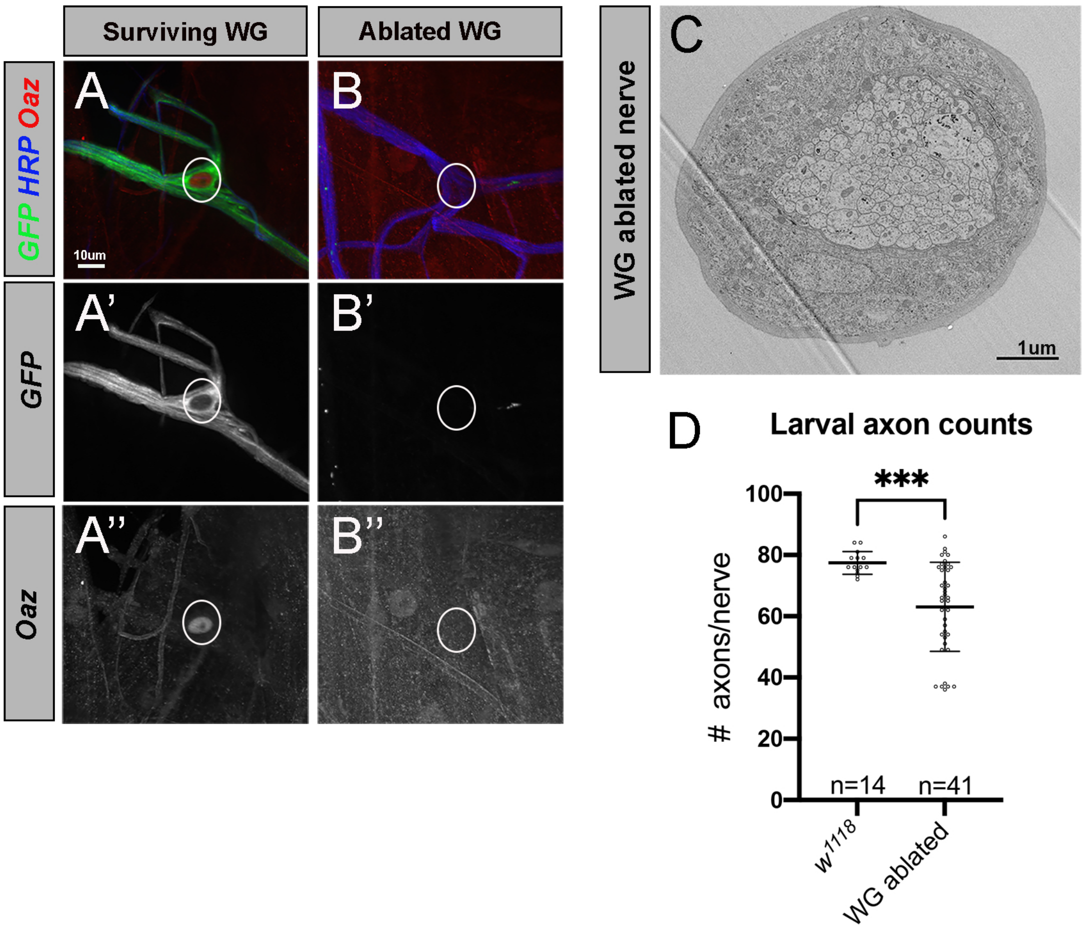

**Supplemental Figure S3: Ablation verification**
Confirmation of ablation using GFP and anti-Oaz staining. A subset of larvae used in behavioral testing were subsequently dissected to confirm successful ablations. The absence of GFP (or presence of only small amounts of GFP+ debris) was observed along nearly all nerves. Some animals were also stained for Oaz, for additional confirmation the wrapping glia were eliminated. Oaz staining was not observed along nerves that did not have any GFP expression (i.e. where glia were ablated). The only Oaz+ nerve nuclei that were observed were in the few GFP+ wrapping glia found that had escaped ablation. **(A)** The cell body region of a surviving ePG5 wrapping glia (von Hilchen et al, 2013) with an Oaz+ nucleus. This surviving cell also continued to express GFP. **(B)** The same stereotyped position along the nerve of the ePG5 cell body in an adjacent segment, but there is no GFP nor Oaz staining, confirming the cell has died rather than just down-regulated GFP. **(C)** TEM cross section of an A8/9 nerve from a wrapping glia ablated animal. There is no observable wrapping glia membrane between any axons, but the outer perineurial glia layer appears hypertrophied. About one quarter of nerves from ablated larvae appeared to show this hypertrophy. **(D)** Axon counts from TEM analyzed A3-A7 larval nerves. Nerves from WG ablated animals frequently have fewer identifiable axon profiles than the wildtype background strain. *($w^{1118}$* average= 77.4 axons; WG-ablated average= 63 axons; Unpaired t-test p=0.0006)

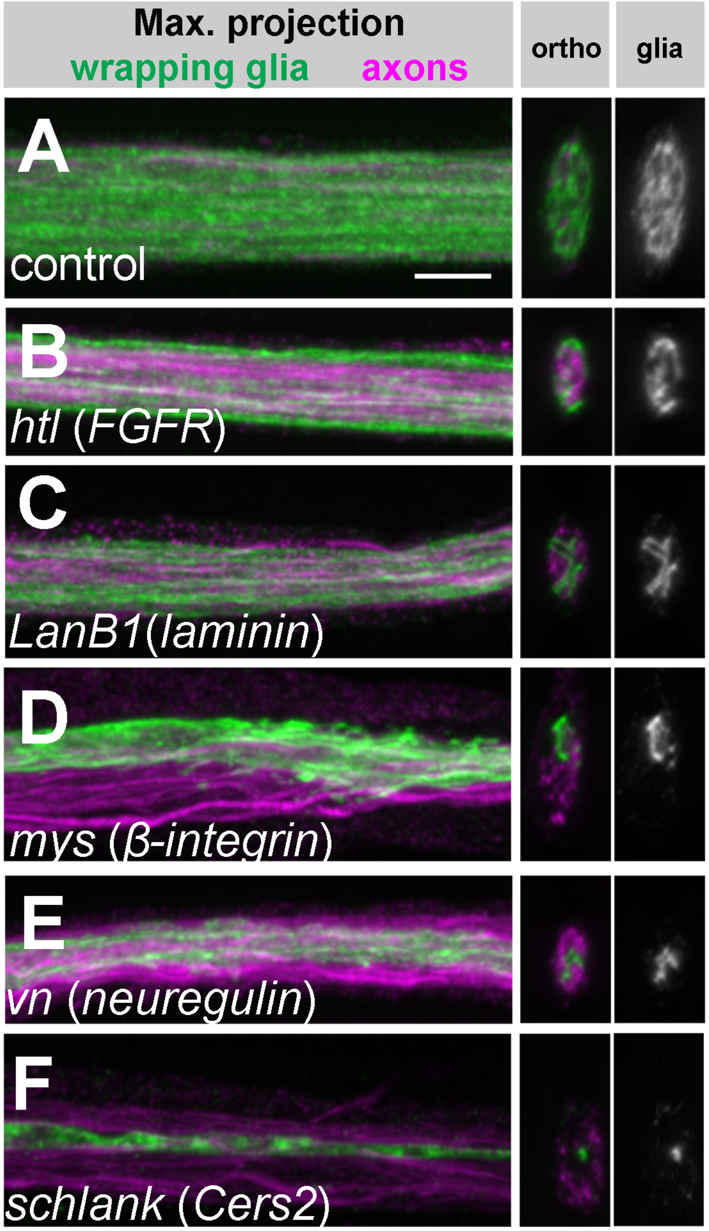

**Supplemental Figure S4:**
**(A-F)** RNAi against genes with known roles in wrapping glia and vertebrate glia development results in visible morphological defects at the light level. Wrapping glia are labeled with myr:tdTomato driven by *nrv2-Gal4* (pseudo-colored green). A subset of sensory axons are labeled with anti-Futsch (magenta). A maximum confocal projection of the nerve is shown in the left panel. The ortho panel shows the nerve cross section, and the glia channel is the isolated wrapping glial channel in cross section. Scale bar 5µm. **(A)** Control nerves show fairly uniform, honeycomb-like glia membrane coverage of the nerve cross section. **(B)**Wrapping glial knockdown of *heartless*, an FGF Receptor homolog, shows incomplete glial coverage of the nerve cross section. **(C)** Knockdown of *lanB1*, a laminin homolog, causes impaired cross section coverage. **(D)** Knockdown of the β-integrin homolog *myospheroid* causes highly abnormal glia morphology where glia fail to cover the entire nerve. **(E)** Knockdown of the neuregulin homolog, *vein*, results in impaired glia coverage of the nerve cross section. **(F)** Knockdown of *schlank*, a Ceramide Synthetase-2 homolog, results in thin stringy glia that does not appear to wrap axons at all.

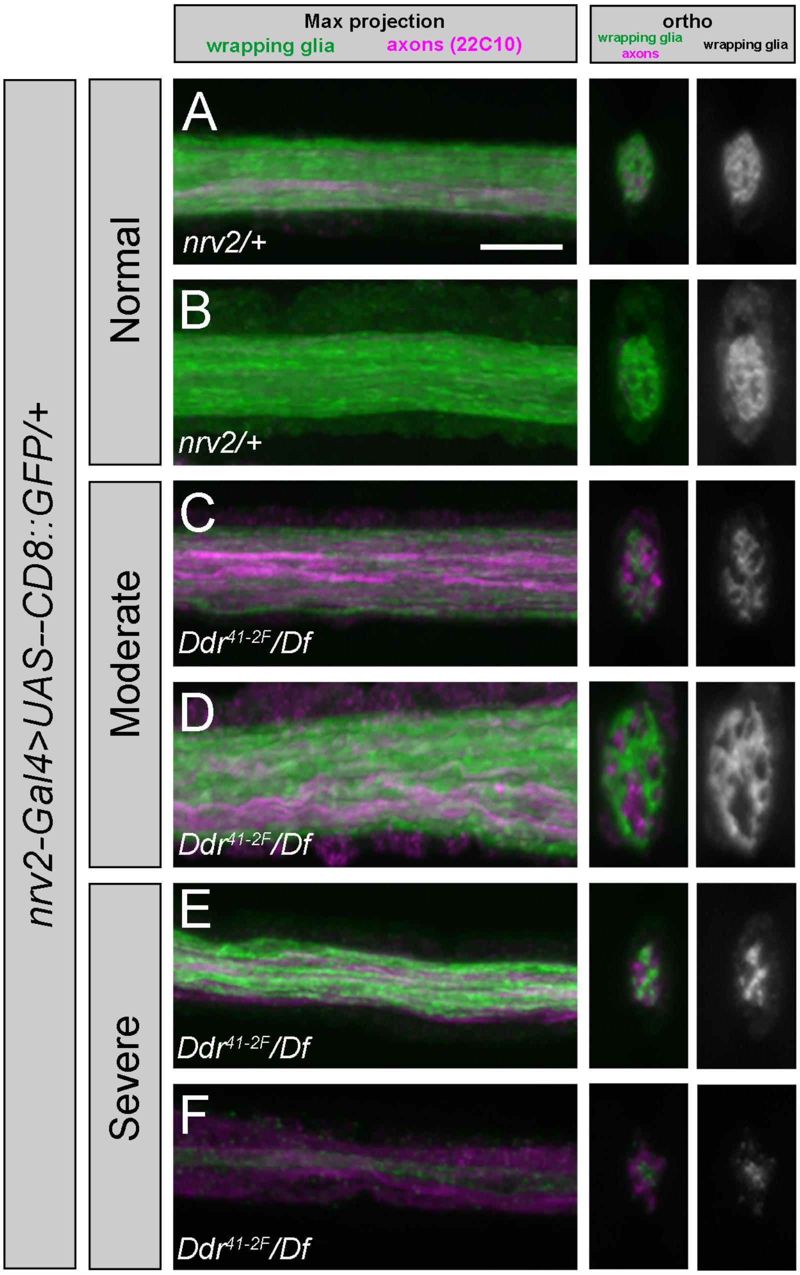

**Supplemental Figure S5: Categorical scoring of wrapping glia morphology**

**(A-B)** Representative images of morphologies classified as "normal" include wrapping glia morphology that displays as a very tight honeycomb (A), to a slightly looser honeycomb with mostly small coverage gaps (B). **(C-D)** Representative images of morphologies classified as "moderate" include wrapping glia with large gaps/openings in coverage of the nerve cross-section though there is some coverage across the entire cross section. **(E-F)** Representative images of morphologies classified as "severe" include wrapping glia that incompletely covers the cross section, often appearing as an "X" type shape in the cross view (E) or a single glial process along and within the nerve (F). Scale bar = 5µm

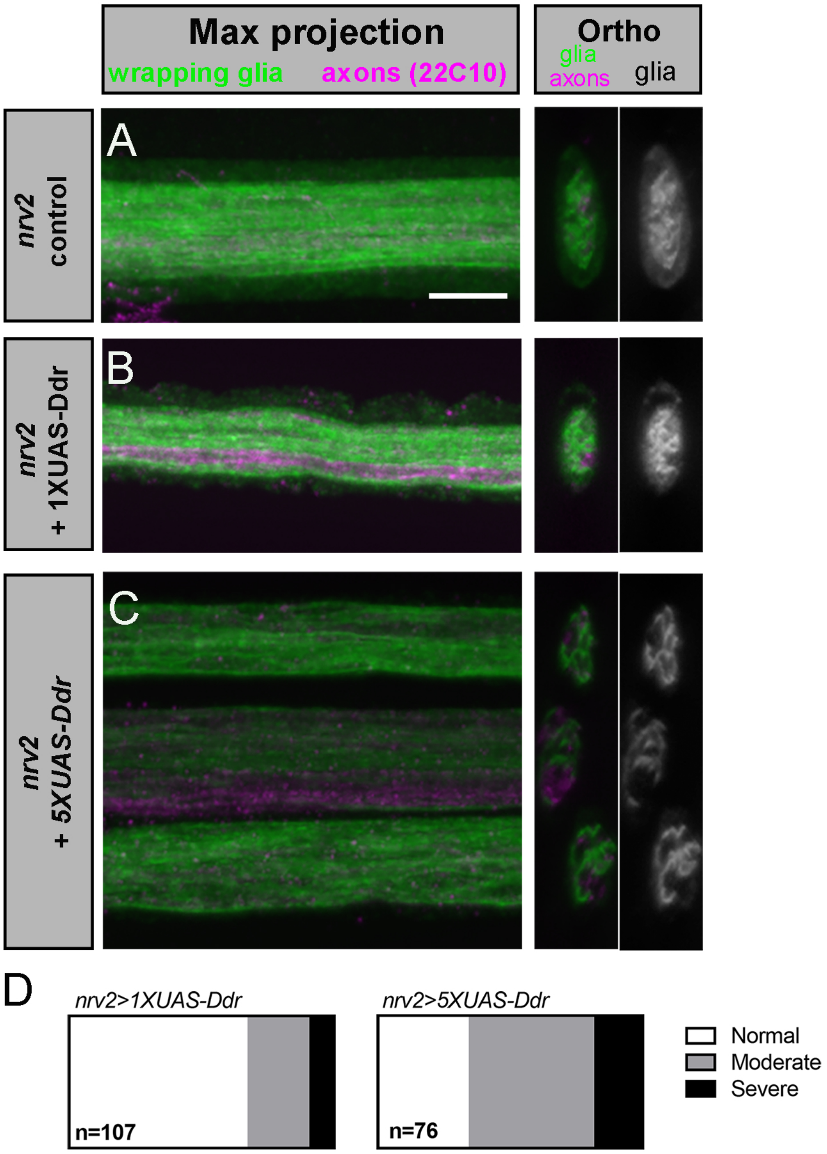

**Supplemental Figure S6: Overexpression of Ddr using a *5XUAS-Ddr* construct causes morphological abnormalities in control animals**

**(A)** *Nrv2-Gal4* driving *UAS-mCD8:GFP* labels wrapping glia showing normal coverage in orthogonal sections. **(B)** Expression of a *1XUAS-Ddr* rescue construct does not cause overexpression artifacts and thus was chosen for mutant rescue experiments. **(C)** A *5XUAS-Ddr* rescue construct frequently caused abnormal wrapping glia morphology. **(D)** Quantification of 1X and 5X constructs in the *nrv2-Gal4* control background. Related to graphs in Figure 2M. n = # of nerves analyzed from 14 (1XUAS) and 10 (5XUAS) larvae.

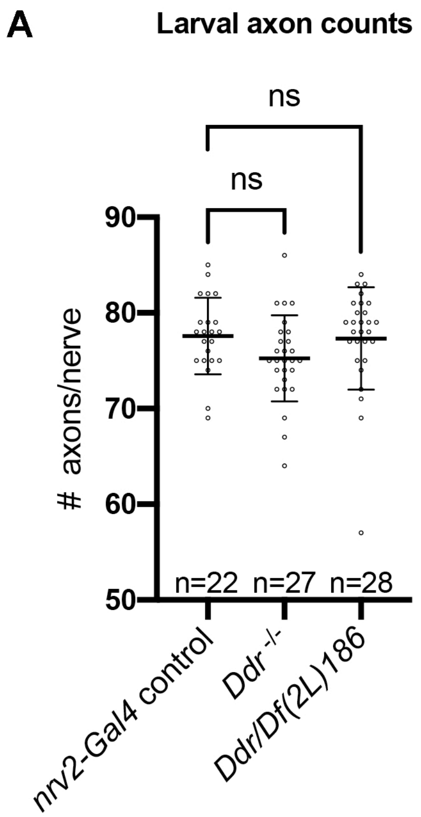

**Supplemental Figure S7: Loss of Ddr does not affect axon profile number**
**(A)** Quantification of axon profiles from A3-A7 nerves in *nrv2-Gal4/+* (control), *Ddr* homozygous mutant, and *Ddr/Df* conditions. There is no significant difference in the number of axon profiles between the conditions. (One-way ANOVA: *Ddr*$^{-/-}$ vs. control p=0.15; *Ddr/Df* vs. control p=0.97, n= # of nerves analyzed.)

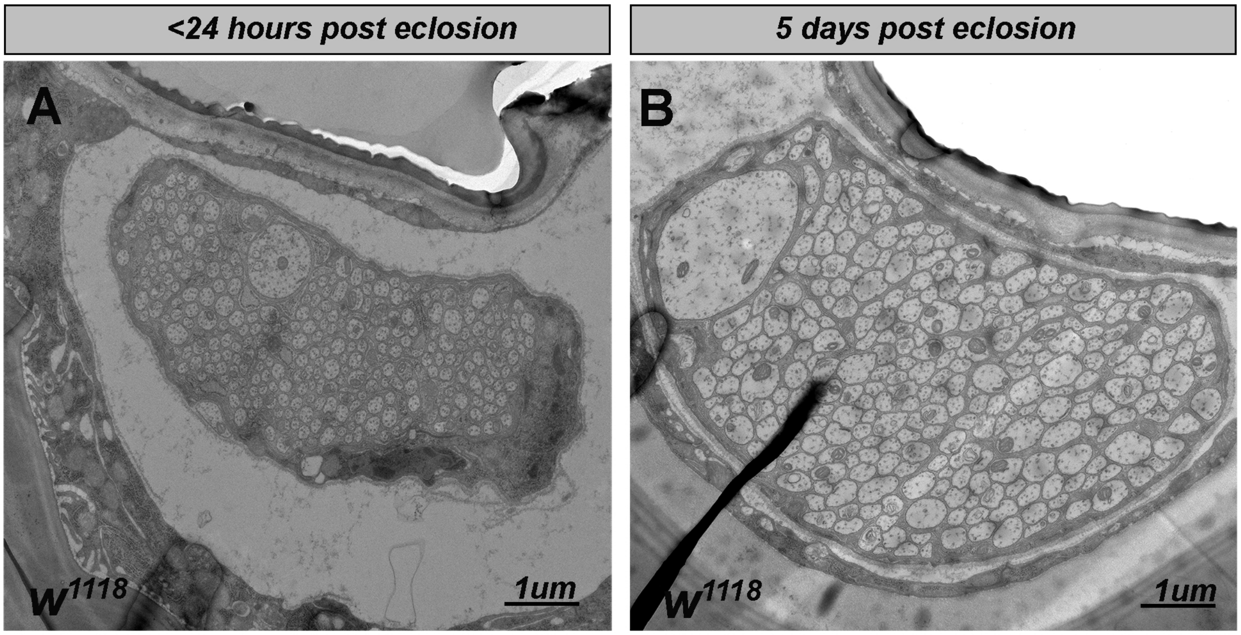

**Supplemental Figure S8: The dTSM axon increases in caliber between eclosion and 5 dpe**

**(A)** Representative TEM of a wild type wing nerve from a female within 24 hours of eclosion shows that the nerve is overall smaller, as is dTSM immediately after eclosion. dTSM is still identifiable as the largest axon in the nerve, but is considerably smaller than in older nerves. **(B)** Representative TEM of a wild type wing nerve from a 5 dpe female.

### Supplemental Methods

#### *Drosophila* stocks

| ***Stock*** | **Source** | **Reference** |
|---|---|---|
| *$w^{1118}$* | | |
| *nrv2-gal4 on 2nd* | Bloomington (BL) #6800 | (Sun et al., 1999) |
| *nrv2-gal4 on 3rd* | BL #6799 | (Sun et al., 1999) |
| *repo-Gal4* | BL #7415 | (Sepp et al., 2001) |
| *nrv2::GFP* | gift of C. Klambt lab | (Stork et al., 2008) |
| *VGlut-QF2* | BL #60315 | (Diao et al., 2015) |
| *OK371(VGlut)-QF2* | BL #66473 | (Lin and Potter, 2016) |
| *UAS-reaper (II)* | BL #5824 | (Aplin and Kaufman, 1997) |
| *IT.0117-Gal4* | BL #62647 | (Gohl et al., 2011) |
| *$IT.0117^{VP16AD}$* | this study | |
| *$nrv2\text{-}Gal4^{DBD}$* | generated in Freeman Lab | (Coutinho-Budd et al., 2017) |
| *$Mp^{f07253}$* | BL #19062 | |
| *$Ddr^{A1\text{-}2F}$* | this study | |
| *$Ddr^{13\text{-}1M}$* | this study | |
| *Df(2L)BSC186* | BL #9614 | (Cook et al., 2012) |
| *1xUAS-Ddr* | this study | |
| *5xUAS-Ddr* | this study | |
| *$CH321\text{-}94A23^{VK31}$* (Ddr BAC) | Genetivision #P3-25 | (Venken et al., 2009) |
| *UAS-CD8::GFP on 2nd* | BL #108068 | (Lee and Luo, 1999) |
| *UAS-CD8::GFP on 3rd* | BL #5130 | (Lee and Luo, 1999) |
| *10xUAS-myr:tdTomato* | BL #32222 | (Pfeiffer et al., 2010) |
| *UAS-mCherrynls* | BL #38424 | |
| *UAS-myrGFP.v5-P2A-H2BmCherry.HA* | gift from J. Dubnau Lab | (Chang et al., 2019) |
| *10xQUAS-6XGFP* | BL #52264 | |
| *$Mp^{MI09316\text{-}GFSTF.0}$* | BL #60567 | |
| *WG-SplitGal4* | this study | |
| *UAS-htlRNAi* | VDRC #27180 | |
| *UAS-LanB1 RNAi* | VDRC #23119 | |
| *UAS-mys-RNAi* | VDRC #103704 | |
| *UAS-vnRNAi* | VDRC #109437 | |
| *UAS-schlankRNAi* | VDRC #109418 | |
| *UAS-Mp RNAi* | VDRC #38188 | |
| *UAS-Ddr-RNAi #1* | VDRC #29720 | |
| *UAS-Ddr-RNAi #2* | VDRC #51719 | |
| *nsyb-Gal4* | BL #51635 | |

New fly strains that were generated for this study were constructed as follows:

*$Ddr^{41\text{-}2F}$* and *$Ddr^{13\text{-}1M}$*: Deletion alleles of *Ddr* were generated using CRISPR-Cas9 gene editing (Port et al., 2014). Briefly, we selected gRNAs targeting the 3rd and 4th exons of *Ddr* and cloned these sequences into pCFD3d under a U6 promoter. *Ddr_ex3_pCFD3d* and *Ddr_ex4_pCFD3d* plasmids were co-injected into *Drosophila* embryos expressing Cas9 in the germline (BL stock 51324; *$w^{1118}$;; PBac{y[+mDint2]=vas-Cas9 3P3-GFP}$^{VK00027}$*; Rainbow Transgenics). Progeny of injected flies were screened for large deletions between the target sites in genomic DNA using primers spanning the predicted deletion that would produce a ~360bp product only if a deletion between gRNA sites had occurred. (Without a large deletion a 11.6kB product would be produced.) Animals harboring large deletions were established as individual stocks by crossing to balancer chromosomes and crossing out *vas-Cas9*. Genomic DNA from each stock was sequenced spanning the predicted deletion to determine if any stocks had frameshifts that introduced early stop codons. *$Ddr^{13\text{-}1M}$* and *$Ddr^{41\text{-}2F}$* and had such frame shifts which resulted in predicted lengths of 150aa and 134aa respectively. As the shorter allele, *$Ddr^{41\text{-}2F}$* was selected for further use in experiments. We frequently performed mutant analysis as *$Ddr^{41\text{-}2F}$ /Df (2L)186* to minimize impact of any unexpected background mutations or suppressors acquired via either the CRISPR strategy or homozygous viable status of the *$Ddr^{41\text{-}2F}$* allele.

gRNA sequences:
- gRNA_Ddr_ex3: GGCCAGTTCGGCCCATG|ATA**TGG**
- gRNA_Ddr_ex4: CCTATGTGATTGAGTAC|TGG**AGG**

Deletion detection primers:
- ddrEx3_F1: CGGAATTCCACTGCTTTGTT
- ddrEx4_R1: CCCAGATGGTTCAGATGGTT

*WG-SplitGal4*: *Nrv2-Gal4$^{DBD}$* was generated as described in (Coutinho-Budd et al., 2017). *IT.0117-Gal4* was converted to *IT.0117-VP16AD* with a series of genetic crosses as described in

Gohl et al, 2011. Once *IT.0117-VP16AD* was established as a stock it was recombined with *Nrv2-Gal4$^{DBD}$* to generate a stable *WG-SplitGal4* stock on chromosome III.

<u>*5XUAS-Ddr* and *1XUAS-Ddr*</u>: A full length 3.165kb Ddr cDNA based on the Ddr-PG isoform sequence listed on Flybase was synthesized and cloned into pUC19 by Genscript USA (Pistacaway, NJ 08854 USA). To construct 5XUAS-Ddr, Ddr was amplified from pUC19-Ddr using primers to introduce a 5' BglII site and Kozak sequence and 3' XhoI site. BglII and XhoI were then used to clone Ddr into the pattB-5xUAS vector. To construct 1xUAS-Ddr, the Ddr sequence was isolated from pattB-5xUAS-Ddr using BglII/XhoI and cloned into the pattB-1xUAS vector. The constructs were injected into $y^1$ $w^{67c23}$; P{CaryP}attP154 embryos (Best Gene) for site-directed intregation onto the 3rd chromosome at position 97D2. Transformants were identified by eye color and stocks were established from single male founders by crossing to balancer females.

Cloning primers:
Forward: GAATACAAGAAGAGAACTCTGAATagatctcaaaacATGCCTGCAATAAAGTTACAAGAATCG
Reverse: gctagcatggtaccataggcctatctcgagTCAATACATGTGTGTATGGGTCTG

**<u>Generation of the anti-Oaz antibody</u>**

Oaz was identified as a potential wrapping glia marker in an enhancer trap screen (M. Freeman, unpublished). Rabbits were immunized with a purified fusion peptide corresponding to the C-terminal 123 amino acids of both predicted Oaz isoforms. The fusion peptide was generated by cloning the corresponding cDNA fragment from clone AT08673 (Berkeley Drosophila Genome Project) into pET28a and inducing protein expression in bacteria. The resulting serum produces optimal staining if a stock of 1:50 is pre-absorbed against either fixed *Drosophila* embryos or fixed 3rd instar larval carcasses overnight at 4°C. This preabsorbed sera can then be diluted 1:100 for a final concentration of 1:5000. Oaz labels nuclei in the CNS (subset of glia and neurons) and muscles in addition to wrapping glia nuclei. Along the nerves, only wrapping glia nuclei are labeled and can thus be positively identified by anti-Oaz staining.

Oaz peptide sequence for immunization:
NHMGEHGAHSRPYDCNLCPEKFFFRAELEHHQRGHELRPQARPPAAKVEVPSIRNTSPGQSPVRSPTIVKQE
LYETDTVESAGVEDEPENHPDEEEYIEVEQMPHETRPSGIGSQLERSTSSA

**Electron Microscopy**

Larvae: 3rd instar larvae were dissected as fillets in ice cold 0.1M cacodylate buffer pH 7.4 (EMS). The buffer was immediately substituted for 2.5% glutaraldehyde in 0.1M cacodylate buffer and left to fix with gentle agitation for 30 minutes at RT, after which dissecting pins were removed and fillets transferred to fresh fixative overnight at 4°C. After 3x 10 minute washes with gentle agitation in 0.1M cacodylate buffer, fillets were incubated in 1% OsO4 (EMS) prepared in ddH2O for 1 hour at RT. After 3x 10 minute washes in ddH2O, fillets were taken through a 4-dilution EtOH dehydration series, with 30%, 50%, 70%, and 95% EtOH for 10 minutes each followed by 2x 10 minute incubation in 100% EtOH, and 2x 10 minute incubation with propylene oxide (PO). Infiltration with 1:1 PO:Epon-812 resin proceeded overnight at RT on a rotating shaker followed by 2 hours infiltration with 100% Epon-812. Larval fillets were flat embedded between 2 sheets of Aclar plastic (EMS) and cured overnight at 60C before being trimmed and re-embedded in coffin molds to prepare for sectioning. 70nm sections were collected from ~200um posterior to the tip of the VNC and placed on 200 mesh copper grids (EMS). Grids were post-stained with 5% uranyl acetate for 20 minutes and Reynolds lead citrate for 8 minutes before being imaged on a Technai T12 electron microscope at 80kV equipped with and AMT digital camera and software.

Wings: To facilitate fixation of the nerve within the wing cuticle we used microwave assisted fixation. Our protocol was adapted from protocols for zebrafish larvae (Cunningham and Monk, 2018; Czopka and Lyons, 2011). Adult flies were anesthetized on CO2 pads and wings were removed with fine dissection scissors and forceps taking care not to touch or crush the anterior edge of the wing. Wings were immediately placed in Eppendorf tubes with modified Karnofsky's fixative (2% glutaraldehyde, 4% paraformaldehyde in 0.1M sodium cacodylate buffer, pH 7.4; EMS). Wings from the same genotype were collected in each tube for primary fixation in a Pelco Biowave microwave (Ted Pella) with the settings: 100W for 1 min, OFF for 1 min x2; 450W for 20 seconds, OFF for 20 seconds x5. Wings were kept in the fixative overnight at 4°C before proceeding. Following 3x10 minute washes with fresh 0.1M sodium cacodylate buffer, 2% OsO4 was added to each tube and samples were again microwaved at 100W for 1 min, OFF for 1 min x2 followed by 450W for 20 seconds, OFF for 20 seconds, x5. OsO4 is then washed out with 3x10minutes washes with ddH20. In bloc UA staining is then performed by adding 8% UA to each tube and microwaving (450W for 1 min, OFF for 1 min, 450W for 1 min). The UA is washed out with 3x10 minute washes with ddH20 and then the dehydration series begins. Samples are taken through an EtOH dilution series of 30%, 50%, 70%, 80%, 95%, each followed by a microwave cycle (250W for 45 seconds) prior to proceeding to the next step. Three changes of 100% EtOH, each followed by a microwave cycle (250W for 1 min, OFF for 1 min, 250W for 1 min) is followed by 3 changes of 100% acetone with the same microwave settings. The final acetone wash is used to move samples to glass dram vials. The pure acetone is replaced with a 1:1 mixture of acetone and Embed 812 (EMS) and allowed to infiltrate samples overnight on a rotating shaker. All microwave steps are carried out with samples in a chilled circulating water bath to keep sample temperature below 20°. Great care must be used when switching solutions as the wings are very hydrophobic and will not sink in liquids until the 80% EtOH step and can be easily lost.

The next day, the 1:1 acetone:Embed-812 mixture is replaced with 100% fresh Embed-812 resin and allowed to infiltrate for at least 1 hour on a rotating shaker. Wings are then flat embedded between Aclar sheets and cured overnight in a 60°C oven. Embedded wings are imaged on Olympus upright microscope to check for any signs of L1 nerve damage. Wings that have tears or scars along the L1 vein are not selected for sectioning and analysis. Intact flat embedded wings are then trimmed close to the ROI with a warm razor blade and re-embedded in coffin molds to facilitate sectioning. 70nm sections are collected on 100mesh formvar film coated grids and counterstained as above with 5% uranyl acetate for 20 minutes and Reynolds lead citrate for 8 minutes before being imaged on a Tecnai T12 electron microscope at 80kV or 120kV equipped with and AMT digital camera and software.

**Full genotypes of animals used in each figure**

| | |
|---|---|
| Figure 1 | |
| A | *$w^{1118}$;;nrv2-Gal4, UAS-mCD8:GFP/+* |
| C | *$w^{1118}$;; nrv2-Gal4, UAS-CD8::GFP/UAS-mCherrynls* |
| D | *$w^{1118}$; ; IT.Gal4-0117/UAS-myrGFP.V5-P2A-H2BmCherry.HA* |
| E | *$w^{1118}$; 5xUAS-mCD8:GFP/+; WG Split- Gal4/UAS-mCherrynls* |
| Figure S1 | |
| A | *$w^{1118}$;; nrv2-Gal4, UAS-CD8::GFP/UAS-mCherrynls* |
| B | *$w^{1118}$; ; IT.Gal4-0117/UAS-myrGFP.V5-P2A-H2BmCherry.HA* |
| C | *$w^{1118}$; 5xUAS-mCD8:GFP/+; WG Split- Gal4/UAS-mCherrynls* |
| Figure 2 | |
| A, D, E | *$w^{1118}$; 5xUAS-mCD8:GFP/+; WG Split- Gal4/+* |
| B, C, D, E | *$w^{1118}$; 5xUAS-mCD8:GFP/UAS-reaper; WG Split- Gal4/+* |
| Figure S2 | |
| A, B | *$w^{1118}$; nrv2-Gal4, UAS-CD8::GFP/UAS-mCherrynls* |

| | |
|---|---|
| C | *$w^{1118}$; 5xUAS-mCD8:GFP/+; WG-Split Gal4/UAS-mCherrynls* |
| Figure S3 | |
| A | *$w^{1118}$; 5xUAS-mCD8:GFP/+; WG-Split Gal4/+* |
| B, C | *$w^{1118}$; 5xUAS-mCD8:GFP/UAS-reaper; WG-Split Gal4/+* |
| Figure 3 | |
| A | *$w^{1118}$; nrv2Gal4, 10XUAS-myrtdTomato/+* |
| B | *$w^{1118}$; nrv2-Gal4, 10XUAS-myrtdtomato/+; UAS-Ddr-RNAi #1* |
| C | *$w^{1118}$; nrv2-Gal4, 10XUAS-myrtdtomato/+; UAS-Ddr-RNAi #2* |
| E | *$w^{1118}$; nrv2Gal4, 5xUAS-mCD8:GFP/+* |
| F | *$w^{1118}$; $Ddr^{13-1M}$/$Ddr^{13-1M}$; nrv2Gal4, 5xUAS-mCD8:GFP/+* |
| G | *$w^{1118}$; $Ddr^{41-2F}$/$Ddr^{41-2F}$; nrv2Gal4, 5xUAS-mCD8:GFP/+* |
| I | *$w^{1118}$; ; nrv2Gal4, 5xUAS-mCD8:GFP/+* |
| J | *$w^{1118}$; $Ddr^{41-2F}$ /Df(2L)BSC186; nrv2Gal4, 5xUAS-mCD8:GFP/+* |
| K | *$w^{1118}$; $Ddr^{41-2F}$ /D()2L)BSC186; nrv2Gal4, 5xUAS-mCD8:GFP/BACP3-25* |
| L | *$w^{1118}$; $Ddr^{41-2F}$ /Df(2L)BSC186; nrv2Gal4, 5xUAS-mCD8:GFP/1xUAS-Ddr* |
| Figure S4 | |
| A | *$w^{1118}$; nrv2Gal4, 10XUAS-myrtdTomato/+* |
| B | *$w^{1118}$; nrv2Gal4, 10XUAS-myrtdTomato/UAS-htlRNAi* |
| C | *$w^{1118}$; nrv2Gal4, 10XUAS-myrtdTomato/UAS-LanB1 RNAi* |
| D | *$w^{1118}$; nrv2Gal4, 10XUAS-myrtdTomato/UAS-mys-RNAi* |
| E | *$w^{1118}$; nrv2Gal4, 10XUAS-myrtdTomato/UAS-vnRNAi* |
| F | *$w^{1118}$; nrv2Gal4, 10XUAS-myrtdTomato/UAS-schlankRNAi* |
| Figure S5 | |
| A, B | *$w^{1118}$; ; nrv2Gal4, 5xUAS-mCD8:GFP/+* |
| C-F | *$w^{1118}$; $Ddr^{41-2F}$/Df(2L)BSC186; nrv2Gal4, 5xUAS-mCD8:GFP/+* |
| Figure S6 | |
| A | *$w^{1118}$; ; nrv2Gal4, 5xUAS-mCD8:GFP/+* |
| B | *$w^{1118}$; ; nrv2Gal4, 5xUAS-mCD8:GFP/1xUAS-Ddr* |
| C | *$w^{1118}$; ; nrv2Gal4, 5xUAS-mCD8:GFP/5xUAS-Ddr* |
| Figure 4 | |
| A | *$w^{1118}$; ; nrv2-Gal4, 5xUAS-mCD8:GFP/+* |
| B | *$w^{1118}$* |
| C | *$w^{1118}$; $Ddr^{41-2F}$ / $Ddr^{41-2F}$; nrv2-Gal4, 5xUAS-mCD8:GFP/+* |
| D | *$w^{1118}$; $Ddr^{41-2F}$ /Df(2L)BSC186 ; nrv2-Gal4, 5xUAS-mCD8:GFP/+* |
| G, H | *$w^{1118}$* |
| G | *$w^{1118}$; ; nrv2-Gal4, 5xUAS-mCD8:GFP/+* |

| | |
|---|---|
| G | $w^{1118}$; Ddr$^{41-2F}$ /+, nrv2-Gal4, UAS-CD8:GFP/+ |
| G | $w^{1118}$; Df(2L)BSC186/+, nrv2-Gal4, UAS-CD8:GFP/+ |
| G, H | $w^{1118}$; Ddr$^{41-2F}$ /Df(2L)BSC186; nrv2Gal4, 5xUAS-mCD8:GFP/+ |
| Figure S7 | |
| A | $w^{1118}$; ; nrv2-Gal4, 5xUAS-mCD8:GFP/+ |
| | $w^{1118}$; Ddr$^{41-2F}$ / Ddr$^{41-2F}$; nrv2-Gal4, 5xUAS-mCD8:GFP/+ |
| | $w^{1118}$; Ddr$^{41-2F}$ /Df(2L)BSC186 ; nrv2-Gal4, 5xUAS-mCD8:GFP/+ |
| Figure 5 | |
| B | $w^{1118}$ |
| C | $w^{1118}$; Ddr$^{41-2F}$ /Df(2L)BSC186; nrv2Gal4, 5xUAS-mCD8:GFP/+ |
| E | $w^{1118}$; TrojanVglut-QF2, QUAS6XGFP; Repo-Gal4/+ |
| | $w^{1118}$; TrojanVglut-QF2, QUAS6XGFP; Repo-Gal4/DdrRNAi51719 |
| F | $w^{1118}$; TrojanVglut-QF2, QUAS6XGFP; Repo-Gal4/+ |
| G | $w^{1118}$; TrojanVglut-QF2, QUAS6XGFP; Repo-Gal4/DdrRNAi51719 |
| I | $w^{1118}$; nrv2Gal4, 5xUAS-mCD8:GFP/+ |
| J | $w^{1118}$; Ddr$^{41-2F}$ /Df(2L)BSC186; nrv2Gal4, 5xUAS-mCD8:GFP/+ |
| Figure 6: | |
| A | $w^{1118}$; TrojanVglut-QF2, QUAS6XGFP; Repo-Gal4/+ |
| B | $w^{1118}$; TrojanVglut-QF2, QUAS6XGFP; Repo-Gal4/DdrRNAi51719 |
| D | $w^{1118}$;;nrv2-Gal4, 5xUAS-mCD8:GFP/+ |
| E | $w^{1118}$; Ddr$^{41-2F}$ /Df(2L) BSC186; nrv2-Gal4, 5XUAS-CD8:GFP/+ |
| G | $w^{1118}$;;nrv2-Gal4, 5xUAS-mCD8:GFP/+ |
| H | $w^{1118}$; Ddr$^{41-2F}$ /Df(2L) BSC186; nrv2-Gal4, 5XUAS-CD8:GFP/+ |
| Figure S8 | $w^{1118}$ |
| Figure 7 | |
| A | $w^{1118}$; nrv2-Gal4, 10XUAS-myrtdTomato/+ |
| B | $w^{1118}$; nrv2-Gal4, 10XUAS-myrtdTomato/UAS-MpRNAi38188 |
| C | $w^{1118}$; Mp MimicGFP/+ |
| D | $w^{1118}$;; Mp$^{f07253}$/nrv2-Gal4, UAS-CD8:GFP |
| E | $w^{1118}$; Ddr$^{41-2F}$ /+, nrv2-Gal4, UAS-CD8:GFP/+ |
| F | $w^{1118}$; Ddr$^{41-2F}$ /+, Mp$^{f07253}$/nrv2-Gal4, UAS-CD8:GFP |
